# Amygdala Reward Neurons Form and Store Fear Extinction Memory

**DOI:** 10.1101/615096

**Authors:** Xiangyu Zhang, Joshua Kim, Susumu Tonegawa

## Abstract

The ability to extinguish conditioned fear memory is critical for adaptive control of fear response, and its impairment is a hallmark of emotional disorders like post-traumatic stress disorder (PTSD). Fear extinction is thought to take place when animals form a new memory that suppresses the original fear memory. However, little is known about the nature and the site of formation and storage of the new extinction memory. Here, we demonstrate that a fear extinction memory engram is formed and stored in a genetically distinct basolateral amygdala (BLA) neuronal population that drive reward behaviors and antagonize the BLA’s original fear neurons. The activation of the fear extinction engram neurons and natural reward-responsive neurons overlap extensively in the BLA. Furthermore, these two neuron subsets are mutually interchangeable in driving reward behaviors and fear extinction behaviors. Thus, fear extinction memory is a newly formed reward memory.

## Introduction

The inability to extinguish fear is a hallmark of many psychiatric disorders, such as PTSD and generalized anxiety disorder (Shalev et al., 2017; Stein and Sareen, 2015). Following Pavlovian fear conditioning (Pavlov and Anrep, 1927), repeated or prolonged presentations of the conditioned stimulus (CS) without an expected aversive unconditioned stimulus (US) diminishes the conditioned fear response, a phenomenon called fear extinction (Herry et al., 2010; Myers and Davis, 2007; Quirk and Mueller, 2008). Fear extinction has been proposed to involve the formation of a new memory in competition with the original fear memory (Bouton, 2004; Quirk and Mueller, 2008; Quirk et al., 2010). The amygdala is a key structure for fear memory (Davis, 1992; Duvarci and Pare, 2014; Ehrlich et al., 2009; Maren and Fanselow, 1996), and is also involved in the fear extinction memory (Amano et al., 2010; Grewe et al., 2017; Herry et al., 2008). However, it is unknown whether the amygdala is the site for the storage of fear extinction memory, and if so, which subset of amygdala neuron stores this memory.

Excitatory neurons in the mouse basolateral amygdala (BLA) respond to both positive and negative valence stimuli (Beyeler et al., 2016; Davis and Whalen, 2001; Kim et al., 2016; Namburi et al., 2015; Redondo et al., 2014), and more than 90% of these neurons are composed of two genetically, functionally and anatomically distinct neuronal populations (Kim et al., 2016; Kim et al., 2017). R-spondin-2-expressing (*Rspo2*^+^) neurons located in the anterior BLA respond to negative valence stimuli and control negative behaviors and memories, whereas protein-phosphatase-1-regulatory-inhibitor-subunit-1B-expressing (*Ppp1r1b*^+^) neurons located in the posterior BLA (pBLA) respond to positive valence stimuli and control appetitive behaviors and memories (Kim et al., 2016). Furthermore, these two neuronal populations antagonize each other through feed-forward inhibitions mediated by local inhibitory interneurons (Kim et al., 2016).

In this study, we investigated a potential role of BLA *Ppp1r1b*^+^ neurons in fear extinction using a contextual fear extinction paradigm. We found that fear extinction memory engram cells are formed and stored within the BLA *Ppp1r1b*^+^ neuronal population, and these engram cells are necessary for suppressing the original fear memory. Furthermore, fear extinction engram cells and natural reward-responsive cells in the pBLA *Ppp1r1b*^+^ neurons are mutually interchangeable in driving reward functions and fear extinction behaviors.

## Results

### BLA *Ppp1r1b*^+^ neurons are activated during contextual fear extinction

Fear extinction phenomena have been observed and studied in both cue-dependent fear conditioning and context-dependent fear conditioning paradigms (Amano et al., 2010; Baldi and Bucherelli, 2014; Herry et al., 2008; Trouche et al., 2013; Zushida et al., 2007). Since the BLA plays a critical role in contextual fear conditioning (Calandreau et al., 2005; Goosens and Maren, 2001; Huff and Rudy, 2004; Redondo et al., 2014) and a substantial amount of information is available regarding the excitatory neuronal subsets in BLA (see Introduction) (Kim et al., 2016), we employed a contextual fear extinction paradigm in this study. On Day 1, the Extinction group received contextual fear conditioning (CFC) in a box where the context served as the CS and three rounds of footshocks served as the US. On Day 2, mice were returned to the conditioned box for 45 min in the absence of footshocks to receive contextual fear extinction training (FET). On Day 3, the mice were tested for 5 min extinction memory retrieval in the conditioned box and then sacrificed (Figure 1A). Two control groups were set up as follows. The Non-Extinction group went through the same protocol as the Extinction group, except that on Day 2 the mice stayed in their home cage (HC) and didn’t receive any extinction training (Figure 1A). The Non-Shock group received the same protocol as the Extinction group, but received no footshocks on Day 1 (Figure 1A). The Extinction group and Non-Extinction group, but not the Non-Shock group, displayed robust freezing behavior on Day 1 that increased as more footshocks were presented (Figure 1B). On Day 3, the Extinction group froze much less than the Non-Extinction group (Figure 1C).

**Figure 1.**
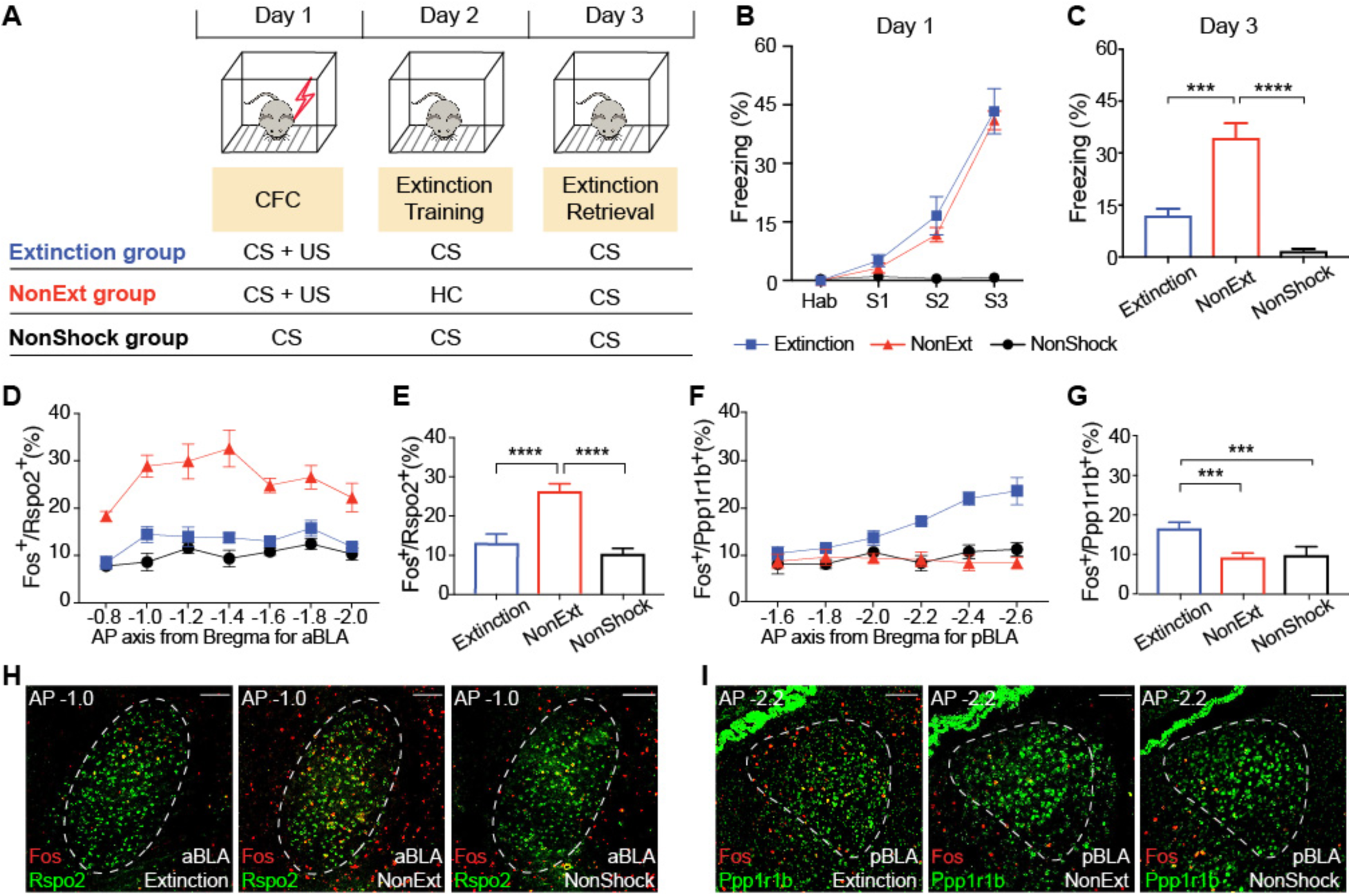
BLA *Ppp1r1b*^+^ neurons are activated during contextual fear extinction. (**A**) Experimental design of the Extinction group, NonExtinction group and NonShock group. (**B** and **C**) Freezing levels of all three groups on Day 1 (B) and Day 3 (C). One-way ANOVA. Extinction *n* = 4; NonExt *n* = 4; NonShock *n* = 4. (**D**) Percentages of *Fos*^+^ neurons within BLA *Rspo2*^+^ neurons across A/P axis, −0.8 mm to −2.0 mm. (**E**) Average of the percentage of *Fos^+^/Rspo2*^+^ shown in (D). One-way ANOVA. Extinction *n* = 4; NonExt *n* = 4; NonShock *n* = 4. (**F**) Percentages of *Fos*^+^ neurons within BLA *Ppp1r1b*^+^ neurons across A/P axis, −1.6 mm to −2.6 mm. (**G**) Average of the percentage of *Fos^+^/Ppp1r1b*^+^ shown in (F). One-way ANOVA. Extinction *n* = 4; NonExt *n* = 4; NonShock *n* = 4. (**H** and **I**) Double smFISH of Fos (red) and Rspo2 (green, H) in aBLA or Ppp1r1b (green, I) in pBLA. \**P* < 0.05, \*\**P* < 0.01, \*\*\**P* < 0.001, \*\*\*\**P* < 0.0001. Data are presented as mean ± SEM. Data are presented as mean ± SEM. Scale bars: 200 µm (H and I)

Double smFISH (single molecular Fluorescence *in situ* Hybridization) was performed to detect the activity-dependent expression of the immediate early gene, *Fos*, in *Rspo2*^+^ and *Ppp1r1b*^+^ cells (see Methods). The proportions of *Fos*^+^ cells of *Rspo2*^+^ cells and *Fos*^+^ cells of *Ppp1r1b*^+^ cells were quantified across the anterior-posterior (A/P) axis for the aBLA and pBLA, respectively (Kim et al., 2016). The proportion of *Fos*^+^/*Rspo2*^+^ was lower in the Extinction group compared to the Non-Extinction group (Figures 1D, 1E and 1H). In contrast, the proportion of *Fos*^+^/*Ppp1r1b*^+^ was higher in the Extinction group compared to the Non-Extinction group while no difference was observed between the Non-Extinction and Non-Shock groups (Figures 1F, 1G and 1I). When BLA *Ppp1r1b*^+^ neurons were optogenetically inhibited during extinction retrieval, the neuronal activity of *Rspo2*^+^ cells (*Fos*^+^/*Rspo2*^+^) as well as the freezing level increased during extinction retrieval (Figures S1). This result suggests that *Ppp1r1b*^+^ neurons suppress *Rspo2*^+^ neurons during fear extinction retrieval, which is consistent with the previously reported feed-forward inhibition of BLA *Rspo2*^+^ cells by BLA *Ppp1r1b*^+^ cells (Kim et al., 2016).

### Dynamics of individual BLA neurons throughout CFC and fear extinction

Next, we performed *in vivo* calcium imaging with a microendoscope to directly track individual BLA neuronal activity during CFC followed by contextual fear extinction. The genetically encoded calcium indicator GCaMP6f was expressed in BLA *Rspo2*^+^ and *Ppp1r1b*^+^ neurons by injecting adeno-associated virus 5 (AAV_5_)-human synapsin (hsyn):double-floxed inverse open reading frame (DIO)*-GCaMP6f* into the aBLA of *Rspo2*-Cre mice and the pBLA of *Ppp1r1b*-Cre mice, respectively (Figures 2A-2C). The efficiency of Cre-loxP recombination in Rspo2-Cre and Ppp1r1b-Cre mice lines are around 90% and 75%, respectively. (Kim et al., 2016). We used an automated sorting algorithm to identify individual neurons and tracked their longitudinal activity across days (Figures 2D, S2A, and S2B; see Methods) (Kitamura et al., 2017; Mukamel et al., 2009; Okuyama et al., 2016). Neuronal activity was analyzed under five conditions: during habituation to the conditioning chamber before CFC (Hab), after footshocks during CFC (Shock), fear retrieval (FR), extinction training (FET) and extinction retrieval (ER) (Figures 2D; see Methods). In order to quantify how neuronal activity changed across these conditions, the activity in each condition was explicitly compared to activity in another condition that served as a baseline, i.e. shock compared to habituation, fear retrieval compared to habituation, FET compared to fear retrieval, and extinction retrieval compared to fear retrieval (Figures 2E and 2F; see Methods). The cells with increased activity were referred to as Up cells, those with decreased activity were referred to as Down cells, and cells whose activity did not change were referred to as No change cells.

**Figure 2.**
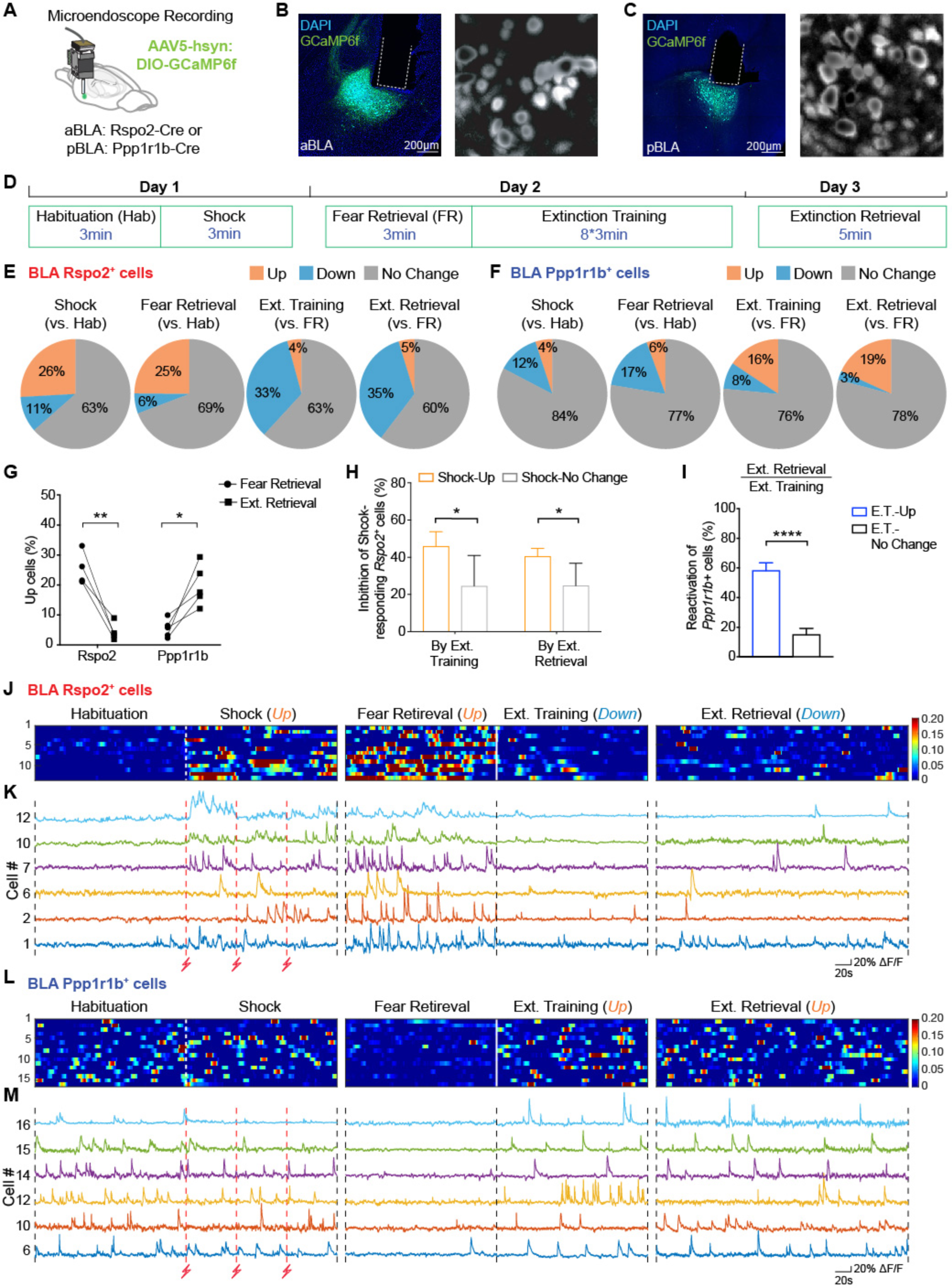
Longitudinal Ca^2+^ imaging of BLA *Rspo2*^+^ and *Ppp1r1b*^+^ neurons during fear extinction. (**A**) Cre-dependent expression of GCamp6f and implantation of microendoscope above the aBLA of Rspo2-Cre and pBLA of Ppp1r1b-Cre mice. (**B**) aBLA of Rspo2-Cre mice. Left: Representative image showing GCamp6f expression and GRIN lens implant; Right: Stacked FOV (Field of View) image. (**C**) pBLA of Ppp1r1b-Cre mice. Left: Representative image showing GCamp6f expression and GRIN lens implant; Right: Stacked FOV image. (**D**) Behavior protocol for Ca^2+^ recording. (**E**) From left to right: percentages of BLA *Rspo2*^+^ neurons with increased (Up, orange), decreased (Down, blue) or No change (grey) responses to shock (vs. habituation), fear retrieval (vs. habituation), extinction training (vs. fear retrieval) and extinction retrieval (vs. fear retrieval). 169 cells total; 4 Rspo2-Cre mice. F.R. represents fear retrieval. (**F**) From left to right: percentages of BLA *Ppp1r1b*^+^ neurons with increased (Up, orange), decreased (Down, blue) or No change (grey) responses to shock (vs. habituation), fear retrieval (vs. habituation), extinction training (vs. fear retrieval) and extinction retrieval (vs. fear retrieval). 179 cells total; 5 Ppp1r1b-Cre mice. F.R.represents fear retrieval. (**G**) Percentages of activated BLA *Rspo2*^+^ and *Ppp1r1b*^+^ neurons during fear retrieval on Day 2 and extinction retrieval on Day 3 per animal. (**H**) Compared to BLA *Rpso2*^+^ neurons with stable responses to footshocks (Shock-No change, grey), BLA *Rpso2*^+^ neurons with increased response to shock (Shock-Up, orange) were preferentially inhibited during extinction training (45.9 ± 3.9% vs. 24.6 ± 8.2%) and extinction retrieval (40.5 ± 2.1% vs. 24.8 ± 12.0%). Paired t-test. (**I**) Compared to BLA *Ppp1r1b*^+^ neurons with stable activity during extinction training (E.T.-No Change), BLA *Ppp1r1b*^+^ neurons with increased activity during extinction training (E.T.-Up) were strongly reactivated during extinction retrieval (58.2 ± 5.3% vs. 15.1 ± 4.2%). Paired t-test. (**J**) Relative fluorescence change (**Δ**F/F) of BLA *Rpso2*^+^ neurons that showed increased activity during shock, increased activity during fear retrieval, decreased activity during extinction training and extinction retrieval (n=13). Each row represents the same cell that was tracked across three days. (**K**) Ca^2+^ traces of six representative BLA *Rspo2*^+^ neurons that are shown in (J) with corresponding cell numbers. (**L**) Relative fluorescence change (**Δ**F/F) of BLA *Ppp1r1b*^+^ neurons that showed increased activity during both extinction training and extinction retrieval (n=16). Each row represents the same cell that was tracked across three days. (**M**) Ca^2+^ traces of six representative BLA *Ppp1r1b*^+^ neurons that are shown in (L) with corresponding cell numbers. \**P* < 0.05, \*\**P* < 0.01, \*\*\**P* < 0.001, \*\*\*\**P* < 0.0001. Data are presented as mean ± SEM.

BLA *Rspo2*^+^ cells were predominantly activated by footshocks during CFC and fear retrieval, and inhibited during extinction training and extinction retrieval (Figures 2E, 2G, and S2C-S2E). Conversely, BLA *Ppp1r1b*^+^ neurons were predominantly activated during extinction training and extinction retrieval, and inhibited by footshocks and fear retrieval (Figures 2F, 2G, and S2F-S2H). A larger percentage of BLA *Rspo2*^+^ cells were activated during fear retrieval compared to extinction retrieval, whereas a larger percentage of BLA *Ppp1r1b*^+^ cells were activated during extinction retrieval compared to fear retrieval (Figure 2G). Compared to BLA *Rspo2*^+^ cells whose activity did not change in response to footshocks (Shock-No Change), shock-activated *Rspo2*^+^ cells (Shock-Up) were preferentially inhibited during extinction training and extinction retrieval (Figures 2H, 2J and 2K). Specifically, 30% of the shock-activated *Rspo2*^+^ cells were also activated during fear retrieval and inhibited during both extinction training and extinction retrieval (Figures 2J and 2K). Compared to BLA *Ppp1r1b*^+^ neurons whose activity did not change by extinction training (E.T.-No Change), *Ppp1r1b*^+^ neurons that were activated during extinction training (E.T.-Up) were preferentially reactivated during extinction retrieval (Figures 2I, 2L and 2M). Combined with the c-Fos quantification data (Figure 1), these results show that pBLA *Ppp1r1b*^+^ neurons are recruited during fear extinction training and reactivated during extinction memory retrieval whereas shock-activated aBLA *Rspo2*^+^ neurons are suppressed during these processes.

### BLA *Ppp1r1b*^+^ neurons drive fear extinction learning

We next investigated the roles of *Ppp1r1b*^+^ and *Rspo2*^+^ neurons in fear extinction at the behavioral level using optogenetic manipulations. We injected a Cre-dependent AAV carrying ChR2 (channelrhodopsin-2) or ArchT (archaerhodopsin) into the aBLA of the *Rspo2-Cre* mice (Figure 3A and S3A) or the pBLA of the *Ppp1r1b*-*Cre* mice (Figure 3B and S3B). Mice underwent the three-day contextual fear extinction protocol as before (Figure 1A). Starting 5 min after the onset of extinction training on Day 2, 3 min of blue or green laser light was delivered during FET repeatedly at 2 min intervals (Figure 3C). Inhibition of *Rspo2*^+^ neurons and activation of *Ppp1r1b*^+^ neurons both facilitated fear extinction learning (Figures 3E and 3F). In contrast, activation of *Rspo2*^+^ neurons and inhibition of *Ppp1r1b*^+^ neurons both impaired fear extinction and extinction memory retrieval (Figures 3D and 3G). These results show that pBLA *Ppp1r1b*^+^ neurons are necessary and sufficient to form fear extinction memory, whereas aBLA *Rspo2*^+^ neurons antagonize it.

**Figure 3.**
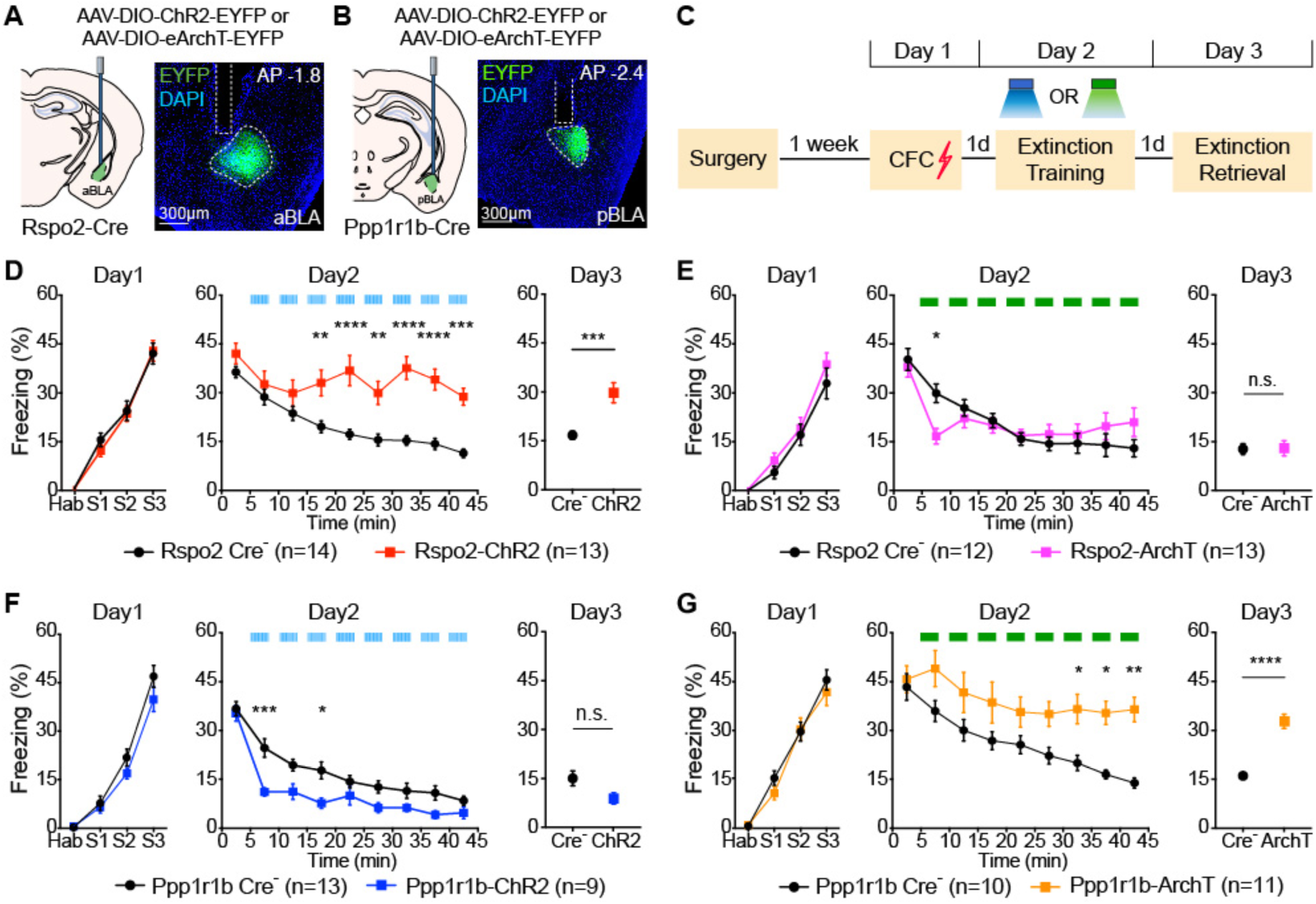
Activation of BLA *Ppp1r1b*^+^ neurons and inhibition of BLA *Rspo2*^+^ neurons facilitate contextual fear extinction. (**A**) Left: diagram of bilateral injection of AAV-DIO-ChR2-EYFP or AAV-DIO-eArchT-EYFP virus and optical fiber implant in the aBLA of Rspo2-Cre mice. Right: representative histology. (**B**) Left: diagram of bilateral injection of AAV-DIO-ChR2-EYFP or AAV-DIO-eArchT-EYFP virus and optical fiber implant in the pBLA of Ppp1r1b-Cre mice. Right: representative histology. (**C**) Experimental protocol of optogenetic manipulation during fear extinction. (**D**) Optogenetic activation of BLA *Rspo2*^+^ neurons impaired fear extinction training and memory. Rspo2-Cre-: *n* = 14, Rspo2-ChR2: *n* = 13. Two-way RM (Repeated Measures) ANOVA. Single-phase exponential decay fit: τ_Rspo2-Cre-_ = 14.06, τ_Rspo2-ChR2_ NA. (**E**) Optogenetic inhibition of BLA *Rspo2*^+^ neurons facilitated early stage of fear extinction training. Rspo2 Cre-: *n* = 12, Rspo2-ArchT: *n* = 13. Two-way RM ANOVA. Single-phase exponential decay fit: τ_Rspo2-Cre-_ = 12.43, τ_Rspo2-ArchT_ = 1.65. (**F**) Optogenetic activation of BLA *Ppp1r1b*^+^ neurons facilitated fear extinction training. Ppp1r1b Cre-: *n* = 13, Ppp1r1b-ChR2: *n* = 9. Two-way RM ANOVA. Single-phase exponential decay fit: τ _Ppp1r1b-Cre-_ = 11.13, τ_Ppp1r1b-ChR2_ = 3.18. (**G**) Optogenetic inhibition of BLA *Ppp1r1b*^+^ neurons impaired fear extinction training and memory. Ppp1r1b Cre-: *n* = 10, Ppp1r1b-ArchT: *n* = 11. Two-way RM ANOVA. Single-phase exponential decay fit: τ _Ppp1r1b-Cre-_ = 11.34, τ_Ppp1r1b-ArchT_ = 20.36. \**P* < 0.05, \*\**P* < 0.01, \*\*\**P* < 0.001, \*\*\*\**P* < 0.0001. Data are presented as mean ± SEM.

### Fear extinction memory engram cells are formed in BLA *Ppp1r1b*^+^ neuronal population

As memories are stored in specific engram cell circuits (Tonegawa et al., 2015), we then investigated whether the engram for fear extinction memory can be identified in the pBLA *Ppp1r1b*^+^ neurons. By using our previously established engram technology (Liu et al., 2012; Ramirez et al., 2013; Redondo et al., 2014; Ryan et al., 2015), we injected AAV_9_-*c-Fos-tTA* (tetracycline transactivator) together with AAV_9_-*TRE* (tetracycline response element)-*ChR2-EYFP* (Ext-ChR2 group) or AAV_9_-*TRE*-*EYFP* (Ext-EYFP group) into the pBLA of C57BL/6 mice (Figures 4A and 4B; see Methods), and subjected them to a series of engram labeling and behavioral steps (Figure 4D; see Methods). This genetic manipulation permits labeling of engram neurons during memory formation or memory retrieval (Figures S4A-S4C) (Khalaf et al., 2018; Liu et al., 2012; Ramirez et al., 2013). During fear extinction training, formation of the putative fear extinction memory engram cells occurs in parallel with retrieval of the original fear memory. In order to isolate the putative fear extinction memory engram neurons, we used the retrieval stage of the fear extinction memory for their labelling (Day3) as fear memory expression will have subsided to the background level by then (Figure 4D).

**Figure 4.**
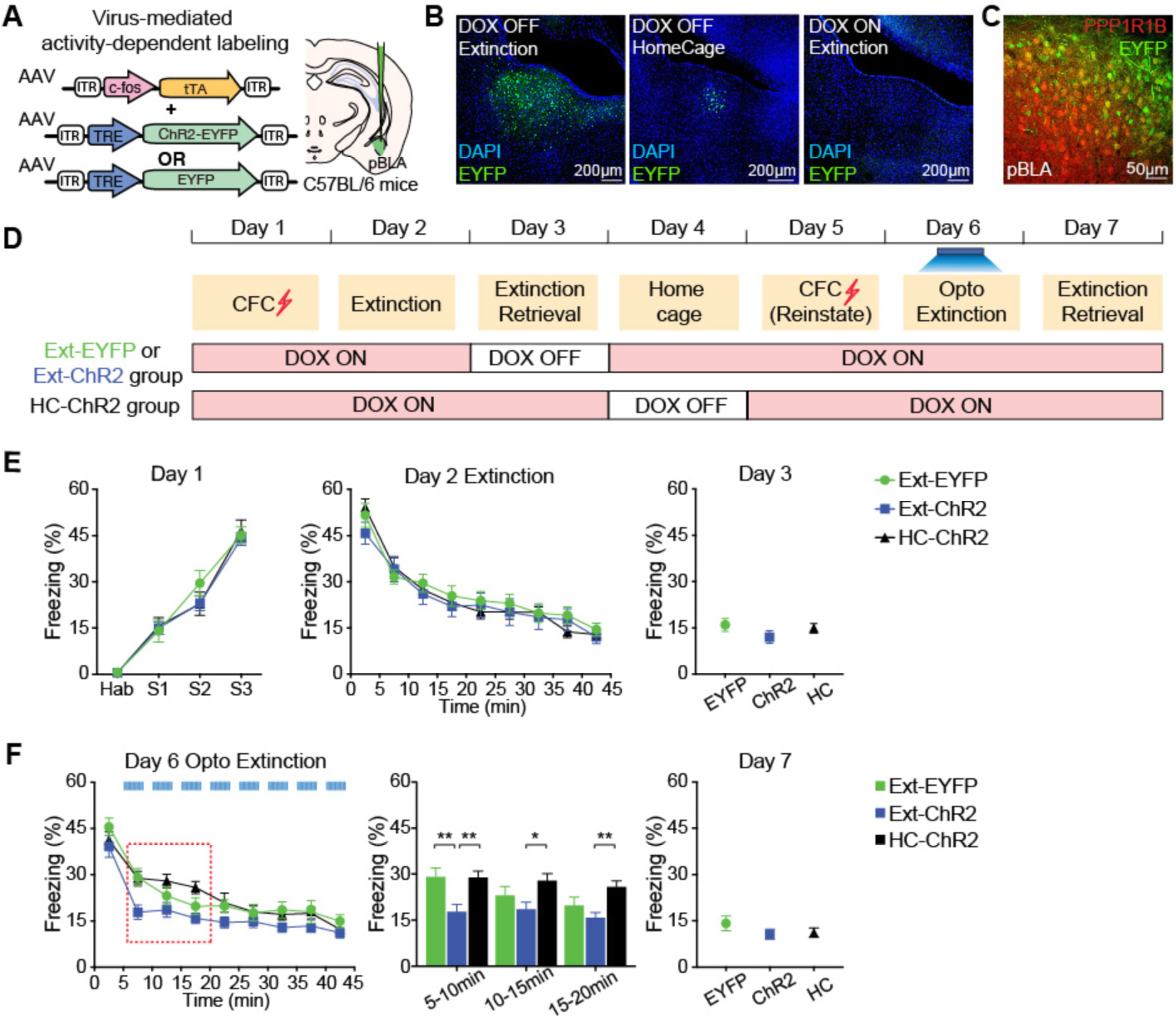
Activation of BLA fear extinction memory engram cells promotes fear extinction. (**A**) Virus-based activity-dependent labeling scheme. C56BL/6 mice were injected with AAV_9_-c-fos-tTA and AAV_9_-TRE-ChR2-EYFP or AAV_9_-TRE-EYFP virus and implanted with optical fibers that bilaterally target pBLA. (**B**) Representative images of EYFP expression under three different conditions: Dox OFF + Extinction (left), Dox OFF + Home Cage (middle) and Dox ON + Extinction (right). (**C**) Image showing 79% of EYFP-expressing cells (green) were PPP1R1B^+^ (red) in pBLA. (**D**) Experimental protocol to label fear extinction engram cells during extinction retrieval in Ext-ChR2 and Ext-EYFP groups. Fear extinction engram neurons were reactivated on Day 6 after reinstatement. In HC-ChR2 control group, ChR2-expressing neurons were labeled when mice were in home cage on Day 4. (**E**) From Day 1 to Day 3, Ext-EYFP, Ext-ChR2 and HC-ChR2 group showed similar freezing level. Two way RM ANOVA test for Day 1 and Day 2. One-way ANOVA test for Day 3. (**F**) On Day 6, reactivation of fear extinction engram cells facilitated fear extinction in Ext-ChR2 group. Bar graph showing freezing levels at 5-10 min, 10-15min and 15-20 min time slots on Day 6 (marked with red dashed line). Two-way RM ANOVA. Ext-EYFP group *n* = 12; Ext-ChR2 group *n* = 16; HC-ChR2 group *n* = 12. Interaction: F (16, 296) = 1.566; P = 0.077. Time: F (8, 296) = 69.65; P < 0.001. Column factor: F (2, 37) = 4.02; P = 0.0263. Subject (matching): F (37, 296) = 7.869; P < 0.001. P (EYPF vs. ChR2) = 0.0020; P (EYPF vs. HC) = 0.9977; P (ChR2 vs. HC) = 0.0025.

A large fraction (79 ± 2%) of pBLA neurons activated by fear extinction retrieval were *Ppp1r1b*^+^ (Figure 4C). In order to access the contribution of food and water supplied in the home cage (HC) to the baseline activity of *Ppp1r1b*^+^ neurons, we set up another negative control group, HC-ChR2 group (Figure 4D). As expected, ChR2-EYFP^+^ cells in the HC-ChR2 group was very sparse compared to those in the ChR2-EYFP group (Figure 4B, S4C-S4D). Three groups of mice, Ext-ChR2 group, Ext-EYFP group and HC-ChR2 group, exhibited indistinguishable freezing levels during the first round of contextual fear extinction protocol from Day 1 to Day 3 (Figure 4E). To examine the roles of these labeled neurons in contextual fear extinction, we subjected the mice to another round of CFC in the same box for fear reinstatement (Day 5). When these neurons were optogenetically activated during the second round of fear extinction training on Day 6 (see Methods), the Ext-ChR2 group showed accelerated extinction compared to the Ext-EYFP and HC-ChR2 groups (Figure 4F). This reactivation-induced accelerated extinction was not observed when the laser was off (Figures S4E-S4F). These results show that the extinction memory engram is formed in the BLA *Ppp1r1b*^+^ neuron population.

### BLA *Ppp1r1b*^+^ extinction engram cells are crucial for suppressing *Rspo2*^+^ fear cells

We then investigated whether pBLA *Ppp1r1b*^+^ extinction engram neurons are not only capable of driving fear extinction, but also necessary for maintaining extinction memory and for suppressing *Rspo2*^+^ fear cells. We labeled the pBLA extinction engram cells with ArchT-mCherry during fear extinction memory retrieval (Figures 5A-5C and S5A). As expected, a large proportion (75 ± 2%) of the pBLA neurons activated by fear extinction retrieval were PPP1R1B^+^ (Figures S5B-S5C). Optogenetic inhibition of these ArchT-labeled neurons specifically impaired the retrieval of fear extinction memory on Day 4 (Figure 5D) and failed to suppress BLA *Rspo2*^+^ fear neurons (Figures 5E-5G). The later is consistent with the feed-forward inhibition of *Rspo2*^+^ neurons by *Ppp1r1b*^+^ extinction neurons within the BLA (Kim et al., 2016). Together, these results demonstrate a causal role of pBLA *Ppp1r1b*^+^ engram cells in extinction memory. These fear extinction engram cells: i) are reactivated during extinction memory retrieval (Figure 2M), ii) have the capacity to drive fear extinction (Figure 4F), and iii) are necessary for suppressing the original fear memory (Figures 5D-5G).

**Figure 5.**
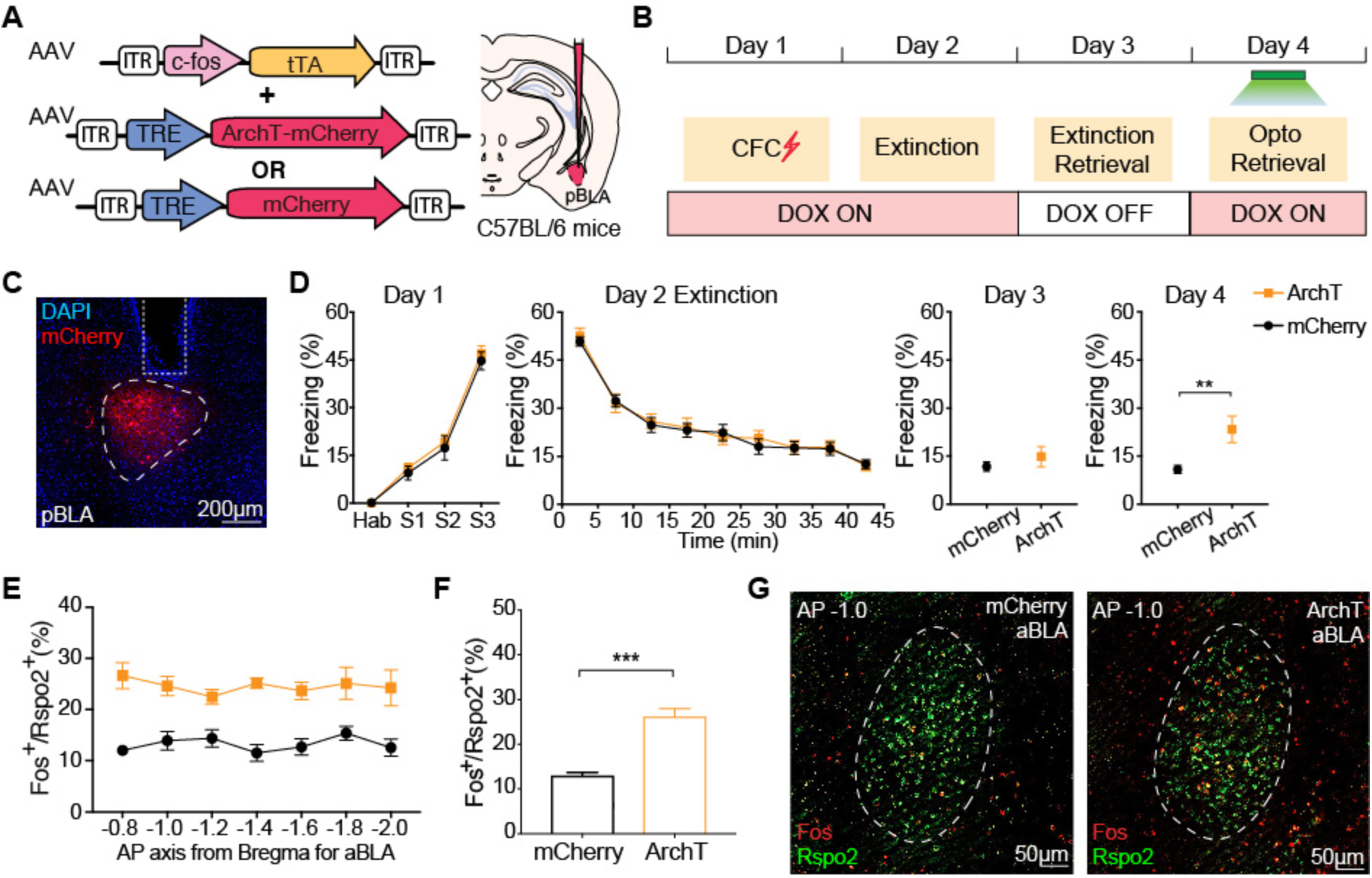
BLA *Ppp1r1b*^+^ fear extinction memory engram cells suppress *Rspo2*^+^ fear cells. (**A**) Virus-based activity-dependent labeling scheme. C56BL/6 mice were injected with AAV_9_-c-fos-tTA and AAV_9_-TRE-ArchT-mCherry or AAV_9_-TRE-mCherry virus and implanted with optical fibers bilaterally targeting pBLA. (**B**) Experimental protocol. Fear extinction engram was labeled on Day 3 during extinction retrieval and inhibited on the following day. (**C**) Representative image showing activity-dependent expression of ArchT-mCherry and optical fiber implant in pBLA. (**D**) Inhibition of fear extinction engram cells in pBLA caused significant recovery of fear response on Day 4. Unpaired t-test. mCherry group *n* = 13; ArchT group *n* = 12. (**E**) When pBLA extinction engram neurons were inhibited during extinction retrieval on Day 4, percentage of *Fos*^+^ neurons within BLA *Rspo2*^+^ neurons across A/P axis, −0.8 mm to −2.0 mm. (**F**) Average of the percentage of *Fos^+^/Rspo2*^+^ shown in (E). Unpaired t-test. Ext-mCherry group *n*=4; Ext-ArchT group *n*=4. (**G**) Double smFISH of Fos (red) and Rspo2 (green) in aBLA of Ext-mCherry mice (left) and Ext-ArchT mice (right). \**P* < 0.05, \*\**P* < 0.01, \*\*\**P* < 0.001, \*\*\*\**P* < 0.0001. Data are presented as mean ± SEM.

### Reward neurons and fear extinction engram neurons overlap extensively in the amygdala

Our previous study demonstrated that pBLA *Ppp1r1b*^+^ neurons responded to a variety of appetitive stimuli (Kim et al., 2016), whereas our present study shows that fear extinction memory is stored in the pBLA *Ppp1r1b*^+^ neuronal population (Figures 1-5). Therefore, we investigated the relationship between these two subsets of *Ppp1r1b*^+^ neurons. To label reward-responding pBLA neurons, AAV_9_*-c-Fos-tTA* and AVV_9_-*TRE-EYFP* virus were injected into pBLA of C57BL/6 mice on Dox-ON diet (Figure 6A). One week post-surgery, all mice were partially deprived of water for 24 hrs on Dox-ON diet (Day 1) and water was given as a reward on Dox-OFF diet (Day 2) (Figure 6B). Mice were then divided into W/W (Water/Water), W/F (Water/Food), W/Ext (Water/Extinction), and W/NonExt (Water/NonExtinction) groups, and kept on Dox-ON diet throughout the remainder of the experiment (Figure 6B; see Methods).

**Figure 6.**
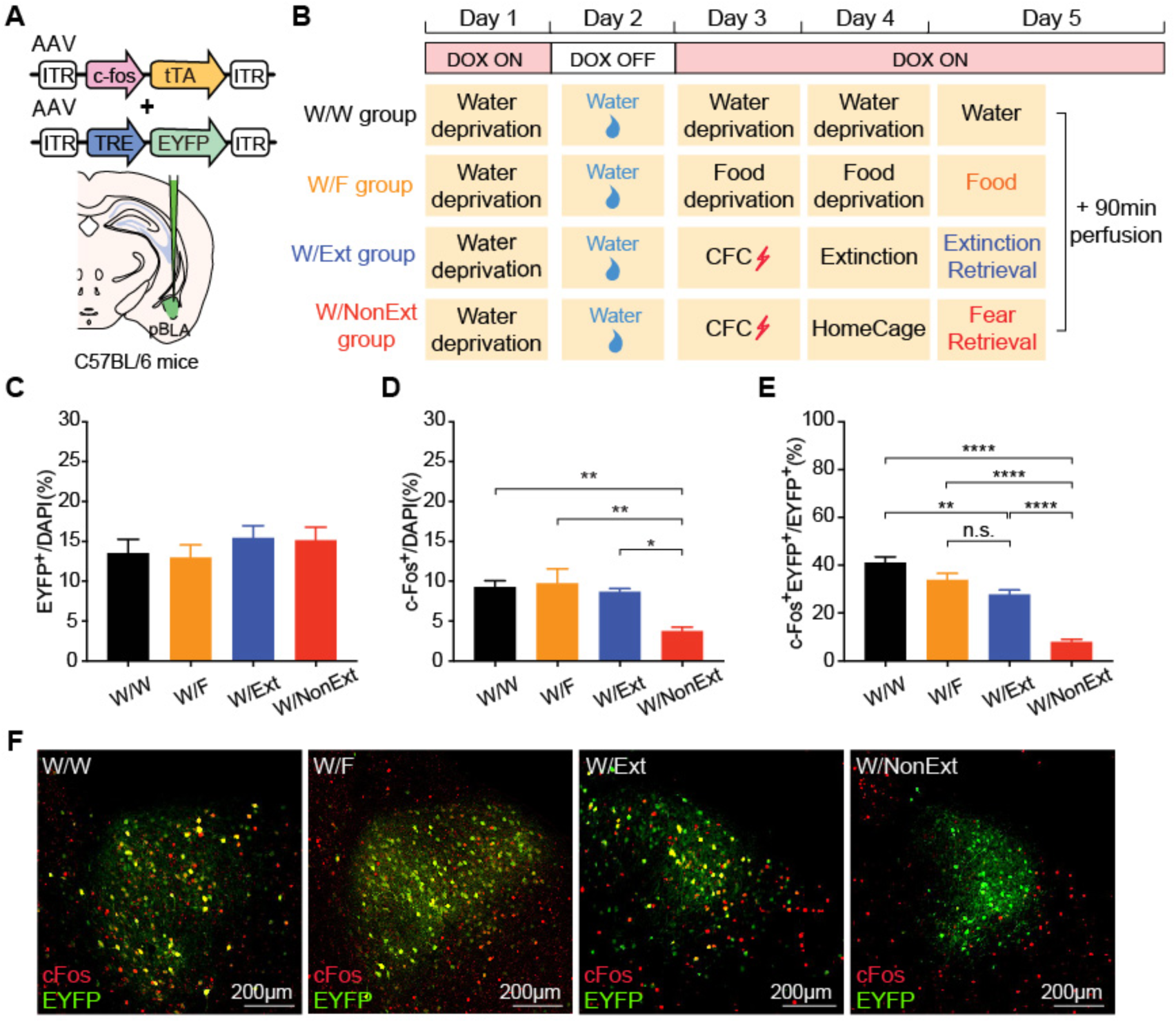
Fear extinction engram neurons and reward neurons overlap extensively on the cellular level. (**A**) Virus-based activity-dependent labeling scheme. C56BL/6 mice were injected with AAV_9_-c-fos-tTA and AAV_9_-TRE-EYFP. (**B**) Experimental protocol for activity-dependent labeling of reward-responsive neurons and detect extinction-activated neurons (W/Ext, blue). Water stimulus (W/W, black), food stimulus (W/F, orange) and fear memory retrieval (W/NonExt, red) were used as controls. (**C**) Similar proportions of neurons were labeled by water reward in all four groups. One-way ANOVA. W/W *n* = 5; W/F *n* = 5; W/Ext *n* = 5; W/NonExt *n* = 5. (**D**) Percentage of c-Fos^+^ cells in pBLA that were activated by water, food, extinction retrieval or fear retrieval. One-way ANOVA. (**E**) Reactivation levels (c-Fos^+^EYFP^+^/EYFP^+^) in W/W group, W/F group and W/Ext group were significantly higher than that in W/NonExt group. No significant difference was observed between W/F group and W/Ext group. One-way ANOVA. (**F**) Double immunofluorescence staining for EYFP (green) and c-Fos (red) in pBLA. \**P* < 0.05, \*\**P* < 0.01, \*\*\*\**P* < 0.0001. Data are presented as mean ± SEM.

Neurons activated by water reward on Day 2 were labeled with EYFP, and their proportions of all neurons (EYFP^+^/DAPI^+^) were similar across all four groups (Figures 6C, 6F and S6A). Neurons activated by the stimulus delivered on Day 5, namely the second delivery of water reward (W/W group), food reward (W/F group), extinction memory retrieval (W/Ext group), and fear memory retrieval (W/NonExt group), were detected by immunohistochemistry for endogenous c-Fos. The proportions of c-Fos^+^ cells of all neurons (c-Fos*^+^/*DAPI^+^) were similar between W/W group, W/F group and W/Ext group, and were substantially higher compared to the W/NonExt group (Figures 6D, 6F, and S6A). The proportion of the c-Fos and EYFP double-positive cells of EYFP^+^ cells (c-Fos^+^ EYFP^+^/EYFP^+^) was used to measure the degree of similarity between the cells responding to the pair of stimuli used in each mouse group. This proportion reached as high as 40% in the W/W group, which represented the ceiling effect of a repeated presentation of the same reward. The proportion of W/Ext group was lower compared to W/W group but it did not differ significantly from the proportion of W/F group, and both of these groups and W/W group gave greater proportions than W/NonExt group (Figures 6E, 6F, and S6A). These results show that the overlap of pBLA *Ppp1r1b*^+^ neurons activated by fear extinction and water reward is as high as the overlap of pBLA *Ppp1r1b*^+^ neurons activated by two *bona fide* rewards (water and food).

### *Ppp1r1b*^+^ fear extinction engram cells and natural reward cells are interchangeable

We then examined whether this observed cellular overlap could also be demonstrated at the behavioral level. To test whether optogenetic activation of fear extinction engram cells can drive appetitive behavior, four groups of C57BL/6 mice were set up (Figures 7A and 7B). In the Ext-ChR2 and Ext-EYFP groups, extinction engram cells were labeled with ChR2-EYFP and EYFP during extinction retrieval on Day 3, respectively (Figures 7B, S7A, and S7B). As controls, neurons activated by water reward and fear memory retrieval were labeled with ChR2 in Water-ChR2 group and CFC-ChR2 group, respectively (Figures 7B, S7C, and S7D). On Day 4, all mice were subjected to an optogenetic self-stimulation behavior test or optogenetic place preference test (Figures 7B, S7E and S7F; see Methods). 50% of Ext-ChR2 group mice and 67% of Water-ChR2 group mice displayed self-stimulation behavior, but not the Ext-EYFP group or CFC-ChR2 group (Figure 7C). In the optogenetic real-time place preference test, mice in the Ext-ChR2 group and Water-ChR2 group spent significantly more time on the stimulated side, whereas Ext-EYFP and CFC-ChR2 groups spent equal times on both sides of the chamber (Figures 7D, 7E, and S7G). Reciprocally, optogenetic activation of ChR2-expressing water reward cells accelerated fear extinction learning (Figures 7F and 7G). Thus, activation of fear extinction engram cells generates appetitive behaviors and activation of reward neurons facilitates fear extinction.

**Figure 7.**
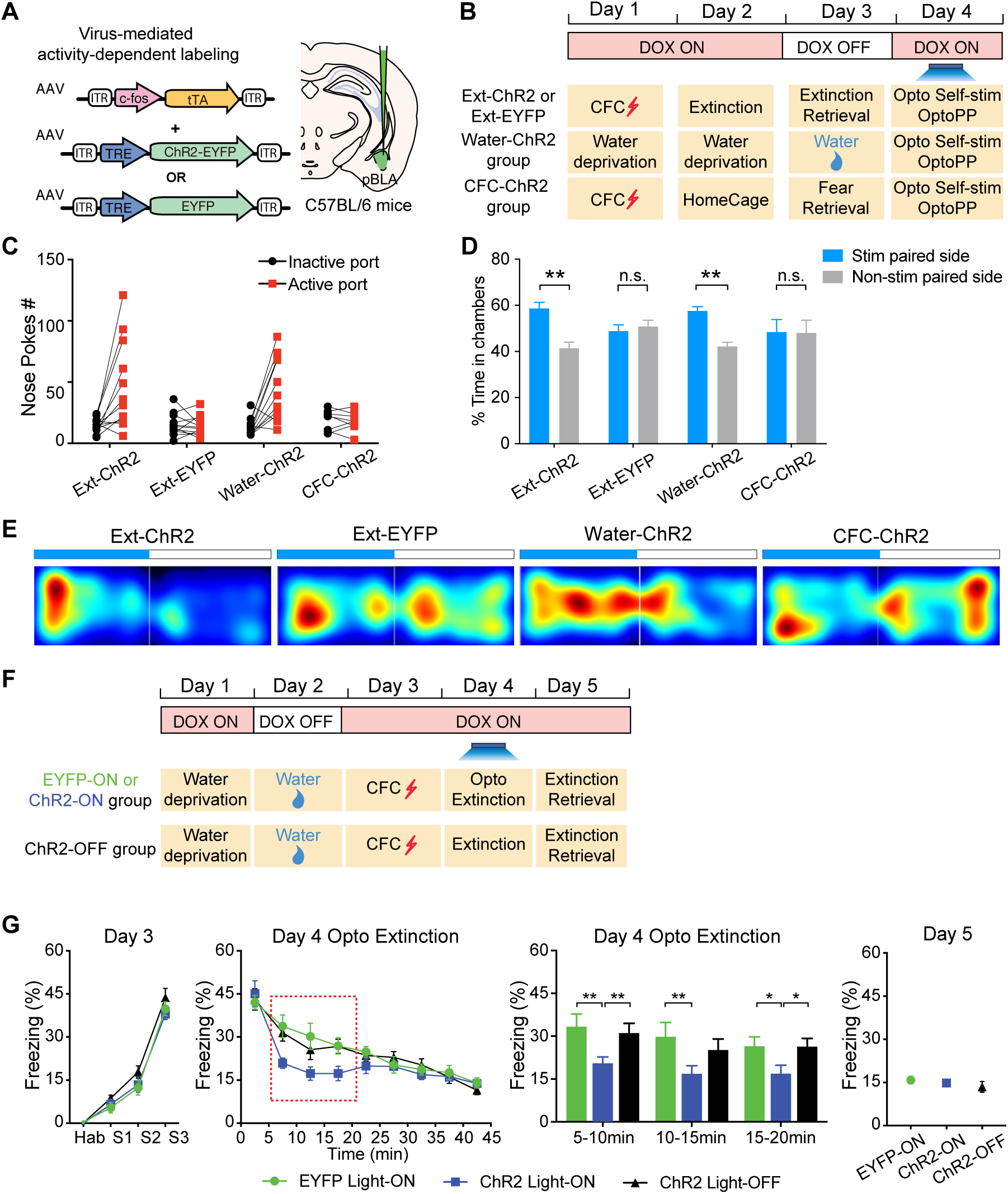
Fear extinction engram neurons and reward neurons are behaviorally interchangeable. (**A**) Virus-based activity-dependent labeling scheme. C56BL/6 mice were injected with AAV_9_-c-fos-tTA and AAV_9_-TRE-ChR2-EYFP or AAV_9_-TRE-EYFP virus and implanted with optical fibers bilaterally targeting pBLA. (**B**) Experimental protocol to label and activate fear extinction engram neurons (Ext-ChR2 or Ext-EYFP group), water-reward neurons (Water-ChR2 group) and fear retrieval engram neurons (CFC-ChR2 group) in optogenetic self-stimulation behavior or optogenetic place preference (OptoPP) behavior on Day 4. (**C**) 50% of Ext-ChR2 group mice and 67% of Water-ChR2 group mice exhibited self-stimulation behavior, with a threshold of (active ports–inactive ports) ≥ 20. No self-stimulation behavior was exhibited in Ext-EYFP or CFC-ChR2 groups. Ext-ChR2 *n* = 12; Ext-EYFP *n* =12; Water-ChR2 *n* = 11; CFC-ChR2 *n* = 8. (**D**) Ext-ChR2 and Water-ChR2 group showed preference towards stimulation paired side. Ext-ChR2 *n* = 12; Ext-EYFP *n* =11; Water-ChR2 *n* = 12; CFC-ChR2 *n* = 11. Paired t-test. (**E**) Heatmaps showing time spent in each side of the chamber. Blue bar showing the stimulation paired side. (**F**) Experimental protocol to activate reward neurons during fear extinction (Day 4). Optical laser was off for ChR2-OFF group during extinction training on Day 4. Optogenetic activation of water-reward neurons accelerated fear extinction learning. EYFP Light-ON, *n* = 8; ChR2 Light-ON, *n* = 11; ChR2 Light-Off, *n* = 10. Two-way RM ANOVA. \**P* < 0.05, \*\**P* < 0.01, \*\*\*\**P* < 0.0001. Data are presented as mean ± SEM.

## Discussion

In this study, we have shown that fear extinction requires new memory engram cells that are formed and stored within BLA *Ppp1r1b*^+^ reward neuronal population. The fear extinction engram cells are defined by their reactivation during extinction retrieval, and their sufficiency and necessity to drive fear extinction. The overlap of neuronal activation of extinction memory engram neurons and natural reward-responsive neurons is extensive and these two types of neurons are mutually interchangeable in driving reward functions and fear extinction behaviors. Thus, the shift from fear to safety in the fear extinction process is mediated by newly formed *Ppp1r1b*^+^ reward memory neurons and their inhibition of the original *Rspo2*^+^ fear memory neurons, both present within the BLA.

The inhibition of *Rspo2*^+^ fear neurons by *Ppp1r1b*^+^ extinction neurons is likely to be mediated by the local inhibitory interneurons in BLA (Kim et al., 2016), supporting the emerging notion that the switch between fear expression and fear extinction takes place within the BLA (Ehrlich et al., 2009; Grewe et al., 2017; Herry et al., 2008; Pare and Duvarci, 2012). Consistent with previous recording studies (Grewe et al., 2017; Herry et al., 2008), our study shows fear extinction is due to new learning rather than a loss of the original fear memory (Herry et al., 2010; Myers and Davis, 2007; Pavlov, 1927). The *Ppp1r1b*^+^ extinction neurons may correspond to the previously reported BLA neurons identified by *in vivo* recording, which were activated throughout extinction learning after cue fear conditioning (Herry et al., 2008). However, our findings extend beyond that study as we further demonstrated that the newly formed extinction memory is an appetitive memory and inhibits the original fear memory.

### Fear extinction recruits the brain’s reward circuit

The finding that the fear extinction memory is a rewarding memory indicates that the emotional valence associated with CS is switched from negative to positive during fear extinction. Notably, the finding that activation of fear extinction engram cells can elicit a reward effect (Figure 7) supports the notion that the absence of expected punishments is in itself a reward. Therefore, fear extinction is positively reinforced by recruiting the brain’s reward circuitry. Previous studies correlated the reward system to fear extinction (Correia et al., 2016; Felsenberg et al., 2018), but their causal relationship was never demonstrated. In contrast, our results show a causal role of the reward system in fear extinction: the pBLA reward neuron population is a necessary component of the fear extinction memory. Furthermore, these findings provide strong evidence of the antagonistic interactions between aversive and appetitive systems that have been observed in Pavlovian fear conditioning (Nasser and McNally, 2012).

### Multiple parallel circuits in fear extinction

In addition to direct feed-forward inhibition within BLA, BLA *Ppp1r1b*^+^ extinction neurons could also drive fear extinction through their interactions with other brain regions. As downstream targets of BLA, both intercalated cells (ITC) and central amygdala (CeA) have been shown to play important roles in fear extinction (Likhtik et al., 2008). Our previous study showed that BLA *Ppp1r1b*^+^ neurons provide monosynaptic projections to the *Prkcd*^+^ neurons located in the lateral subnuclei of CeA (CeL), which are also activated during fear extinction retrieval (Kim et al., 2017). Therefore, the fear extinction behavior can also be modulated via BLA *Ppp1r1b*^+^-CeA monosynaptic projection or BLA *Ppp1r1b*^+^-ITC-CeA di-synaptic projection. The infralimbic region (IL) of the prefrontal cortex is another critical cite for fear extinction (Burgos-Robles et al., 2007; Laurent and Westbrook, 2009; Vidal-Gonzalez et al., 2006) and its connections with BLA *Ppp1r1b*^+^ neurons may also be involved in this process (Cho et al., 2013; Kim et al., 2016; Likhtik et al., 2005; Quirk et al., 2006; Sotres-Bayon et al., 2012).

### Intrinsic reinforcement signals during fear extinction learning

How *Ppp1r1b*^+^ cells are activated for the acquisition of fear extinction memory is another question that remains to be elucidated. Reinforcement learning theories would predict that acquisition of extinction memory is instructed by negative prediction error, in which the predictive value of the CS with respect to the occurrence of the US is compromised (McNally et al., 2011; Schultz, 2016). Given there is no obvious external US in the case of fear extinction, this stimulus may be provided internally and preferentially to *Ppp1r1b* neurons in the BLA. The identification of the agent and the circuit that provide this stimulus would be of fundamental importance.

Accumulating evidence suggests that dopamine reinforces fear extinction learning by signaling the omission of expected aversive US in both animals and humans (de Jong et al., 2019; Kalisch et al., 2019; Matsumoto and Hikosaka, 2009; Raczka et al., 2011; Ray Luo, 2018; Salinas-Hernandez et al., 2018). In the mesolimbic dopaminergic system, amygdala receives dopaminergic inputs from the ventral tegmental area (VTA) and substantia nigra (SNc) in the midbrain (Brinley-Reed and McDonald, 1999; Pezze and Feldon, 2004). Future studies are needed to investigate how dopamine functions as an instructive signal onto BLA neurons for fear extinction learning.

In conclusion, our study provides the empirical evidence for the key concept derived from learning theory that an omission of the expected punishment is rewarding(Felsenberg et al., 2018; Kim et al., 2006; Nasser and McNally, 2012), pinpointing potential targets for effective interventions of fear-related disorders (Shalev et al., 2017; Stein and Sareen, 2015).

## Acknowledgments

We thank X. Zhou, K. Marshall, B. Bane, F.J. Bushard and W. Yu for technical assistance; C. Sun and C.J. MacDonald for discussion on the calcium imaging data analysis; M. Pignatelli, S. Muralidhar, C. Sun and C.J. MacDonald for comments and discussion on the manuscript; and all members of the Tonegawa laboratory for their support.

## Author contributions

X.Z., J.K. and S.T. contributed to the study design. X.Z. contributed to the data collection and interpretation. X.Z. conducted the surgeries, behavior experiments and histological analyses. X.Z. and S.T. wrote the paper. All authors discussed and commented on the manuscript.

## Data and material availability

AAV_9_-c-Fos-tTA, AAV_9_-TRE-ChR2-EYFP, AAV_9_-TRE-eArchT-mCherry, and AAV_9_-TRE-EYFP were developed at the Massachusetts Institute of Technology by the group of S.T.; virus plasmids are available through a material transfer agreement. The Rspo2-Cre transgenic mouse line was developed at RIKEN BSI by the group of Shigeyoshi Itohara and is available through a material transfer agreement. All data is available in the main text or the supplementary materials.

## Supplementary Materials

### Materials and Methods

#### Subjects

The Ppp1r1b-Cre (also called Cartpt-Cre) mice were obtained from GENSAT (stock number: 036659-UCD). The Rspo2-Cre mice were generated in house. Cre-expressing mice were genetic knock-in mice or have been previously been validated for genetic specificity (*16*). All Cre transgenic mice were bred using a heterozygous male with females of C57BL/6 background and maintained as heterozygous. Mice had access to food and water *ad libitum* and were socially housed in numbers of two to five littermates until surgery. After surgery, mice were singly housed. 2-to 6-month old male mice were used for all experiments. For engram labeling, C57/B6 mice were raised on food containing 40 mg kg^−1^ DOX 1 week before stereotaxic surgery and were maintained on Dox diet for the remaining experiment period except for the target Dox-OFF days. All mice were maintained and cared in accordance with protocols approved by the Massachusetts Institute of Technology (MIT) Committee on Animal Care (CAC) and guidelines by the National Institutes of Health (NIH).

#### Viral constructs

AAV_5_-Ef1a-DIO-eArch3.0-EYFP (AV5257), AAV_5_-Ef1a-DIO-ChR2-EYFP (AV5226B), and AAV_5_-Ef1a-DIO-eYFP (AV4310D) were obtained from the University of North Carolina at Chapel Hill Vector Core. AAV_9_-c-fos-tTA, AAV_9_-TRE-EYFP, AAV_9_-TRE-ChR2-EYFP, AAV_9_-TRE-mCherry, and AAV_9_-TRE-eArchT-mCherry were generated by and acquired from the Gene Therapy Center and Vector Core at the University of Massachusetts Medical School as previously described (Kitamura et al., 2017; Liu et al., 2012; Redondo et al., 2014). The AAV_5_-hsyn:DIO-GCaMP6f virus was generated by and acquired from the University of Pennsylvania Vector Core.

#### Stereotactic surgery

Mice undergoing stereotactic injections were anesthetized under isoflurane. Standard stereotactic procedures were used. Viruses were injected using a mineral oil–filled glass micropipette attached to a 1 µl Hamilton microsyringe. Virus was injected at a speed of 80 nl/min, which was controlled by a microsyringe pump. The needle was lowered to the target site and remained for 5 min after the injection. The incision was closed with sutures. Mice were given 0.1 mg/kg slow release Buprenex as analgesic and remained on a heating pad until full recovery from anesthesia. The histology of all injections was verified after behavior experiments.

For optogenetic behavior experiment, 200 µL of AAV_5_ Cre-dependent virus was bilaterally injected into the anterior BLA of Rspo2-Cre mice (AP −1.4 mm, ML ±3.4 mm, DV −4.9 mm) and the posterior BLA of Ppp1r1b-Cre mice (AP −2.0 mm, ML ±3.4 mm, DV −4.9 mm) and incubated for 3 – 4 weeks before behavioral experiments.

For activity-dependent labeling and activation/inhibition of pBLA engram cells, 200 nL of ∼2.0 × 10^9^ GC of AAV_9_-cfos-tTA and AAV_9_-TRE-ChR2-EYFP or AAV_9_-TRE-eArch3.0-mCherry (1:1 mixture) were bilaterally injected into the pBLA (AP −2.0 mm, ML ±3.4 mm, DV −4.9 mm) of doxycycline (Dox)-fed C57/B6 mice and incubated for 7 days before subsequent experiments. AAV_9_-TRE-EYFP or AAV_9_-TRE-mCherry was used as corresponding controls.

For optical implant surgery, two optic fibers (200 mm core diameter, 5.5 mm length; Doric Lenses) were bilaterally lowered above the injection sites (aBLA: −1.4 mm AP, ±3.4 mm ML, −4.7 mm DV; pBLA: −2.0 mm AP, ± 3.4 mm ML, −4.7 mm DV). The implants were secured to the skull with two jewelry screws and adhesive cement (C&B Metabond). A protective cap, made using a 1.5 mL black Eppendorf tube, was fixed onto the implant using dental cement (Kim et al., 2016; Kim et al., 2017).

For Ca^2+^ imaging surgery, AAV_5_-hsyn:DIO-GCaMP6f was unilaterally injected into right aBLA of Rspo2-Cre mice and right pBLA of Ppp1r1b-Cre mice with the following coordinates: aBLA was targeted at −1.4 mm AP, +3.4 mm ML, −4.9 mm DV and pBLA was targeted at −2.0 mm AP, +3.4 mm ML, −4.9 mm DV. Two weeks following AAV_5_-hsyn:DIO-GCaMP6f virus injection, a GRIN lens (0.5 mm diameter, 8 mm length; Inscopix) was implanted targeting aBLA or pBLA (aBLA: −1.4 mm AP, +3.4 mm ML, −4.7 mm DV; pBLA: −2.0 mm AP, +3.4 mm ML, −4.7 mm DV).

#### Contextual fear extinction behavior protocol

Behavioral context was 29 × 25 × 22 cm chambers (Med Associates) with grid floors, opaque ceilings, white lighting, and scented with 5% benzaldehyde. Before fear conditioning, mice were habituated to investigator handling for 5 min on three consecutive days in the holding room where the mice were housed. On Day 1, mice were habituated in the behavioral context for 3 min, followed by three footshocks (0.60 mA, 2 s) delivered at 180 s, 250 s and 320 s. Mice remained in the behavior chamber for 80 s after the third footshock and then returned to their home cages in the holding room. 24 hrs after fear conditioning, mice returned to the fear conditioning chambers to receive 45 min extinction training without any footshocks. Then mice were put back to their home cages for another 24 hrs. On Day 3, mice were placed in the conditioned context for 5 min extinction retrieval test. Behavior videos were recorded with VideoFreeze software and freezing level was scored manually by experimenters who were blinded to conditions.

#### Single molecular fluorescent *in situ* hybridization

To examine the expression of *Fos* gene (Figures 1 and 5), single molecule fluorescence *in situ* hybridization (smFISH) was performed using RNAscope Fluorescent Multiplex Kit (Advanced Cell Diagnostics, ACDBio) as previously described (*16, 18*). Isoflurane anesthetized mice were decapitated, their brains harvested and flash frozen on aluminum foil on dry ice. Brains were stored at −80 °C. Prior to sectioning, brains were equilibrated to −16 °C in a cryostat for 30 min. Brains were coronally sectioned at 20 µm with cryostat and thaw-mounted onto Superfrost Plus slides (25 × 75 mm, Fisherbrand). Sections from a single brain were serially thaw-mounted onto 10 slides through the entire BLA (anterior-posterior distance from Bregma, −0.8 mm to −2.6 mm). Slides were air-dried for 60 min at room temperature prior to storage at −80 °C. smFISH probes for all genes examined were obtained from ACDBio, *Fos* (Cat #421981), *Ppp1r1b* (Cat #405901) and Rspo2 (Cat #402001). Slides were counterstained for the nuclear marker DAPI using ProLong Diamond Antifade mounting medium with DAPI (ThermosFisher) (Kim et al., 2016; Kim et al., 2017).

#### Immunohistochemistry

Mice were perfused transcardially with phosphate buffered saline (PBS), followed by 4% paraformaldehyde (PFA). Brains were post-fixed with 4% PFA solution overnight and then sectioned coronally 50 µm at with a vibratome. Free floating brain sections were washed in PBST (PBS, 3% TritonX) three times for 10 min, blocked for 1 hr in blocking buffer (PBST, 5% normal goat serum), and incubated in primary antibody in blocking buffer overnight at 4 °C. On the next day, brains were washed with three 10 min washes of PBST and incubated in secondary antibody in blocking buffer at room temperature for 2 hrs. Primary antibodies used were chicken anti-GFP (Invitrogen, A10262, 1:1,000), rabbit anti-c-Fos (Santa Cruz, sc-52, 1:1,000) and rabbit anti-PPP1R1B (Abcam, ab40801, 1:1,000). Secondary antibodies used were goat anti-chicken Alexa Fluor 488 (Invitrogen, A11039, 1:1,000) and goat anti-rabbit Alexa Fluor 555 (Invitrogen, A21428, 1: 1,000). After three more 10-min PBST washes, slides were covered with coverslips and mounted using VectaShield mounting solution containing DAPI (Vector Laboratories).

#### Calcium imaging

Calcium imaging of BLA *Rspo2*^+^ neurons and *Ppp1r1b*^+^ neurons was performed on Rspo2-Cre and Ppp1r1b-Cre mice, respectively. AAV_5_-hsyn:DIO-GCaMP6f was unilaterally injected into right aBLA of Rspo2-Cre mice and right pBLA of Ppp1r1b-Cre mice with the following coordinates: aBLA was targeted at −1.4 mm AP, +3.4 mm ML, −4.9 mm DV and pBLA was targeted at −2.0 mm AP, +3.4 mm ML, −4.9 mm DV. Two weeks following AAV_5_-hsyn:DIO-GCaMP6f virus injection, a GRIN lens (0.5 mm diameter, 8 mm length; Inscopix) was implanted targeting aBLA or pBLA (aBLA: −1.4 mm AP, +3.4 mm ML, −4.7 mm DV; pBLA: −2.0 mm AP, +3.4 mm ML, −4.7 mm DV). Two weeks after GRIN lens implant, a baseplate for the miniaturized microscope camera was attached above the GRIN lens using ultraviolet-light curable glue (Loctite 4305) (Mukamel et al., 2009; Ziv et al., 2013).

For contextual fear extinction behavior, recording was performed during the light cycle. Inscopix endoscope and calcium events were captured at 20 Hz with an Inscopix miniature microscope (nVista HD). Inscopix endoscope was synchronized with Med Associate behavior chamber via a TTL signal, which was used for all the other fear extinction behavior experiments in this study. Mice underwent the three-day contextual fear extinction procedure as previously described (Figure 1). During contextual fear conditioning (CFC) on Day 1, freezing behavior and Ca^2+^ signals of subjects were recorded for 3 min habituation and 3 min when three footshocks were delivered. During 45 min fear extinction training, Ca^2+^ signals was recorded during the 0-3 min, 5-8 min, 10-13 min, 15-18 min, 20-23 min, 25-28 min, 30-33 min, 35-38 min and 40-43 min timestamps. The 0-3 min epoch was considered as fear retrieval stage. The other eight epochs together were considered as extinction training stage. In the heatmap and Ca^2+^ traces (Figure 2E-2I), extinction training referred referred to 40-43 min time bin recording on Day 2. Recorded calcium-imaging movies were first processed with ImageJ to subtract base line activity (dividing each image by a low-pass, r = 20 pixels, and high pass, r = 1000 pixels), followed by motion correction using Inscopix Mosaic software (Mukamel et al., 2009). Subsequently, ΔF/F ((F-F0)/F0) signal was calculated in Mosaic software (Inscopix), in which F0 was calculated over the entire recording session. A stacked image of ΔF/F recording movie was acquired by maximum intensity projection. Cell identifications were semi-automatically identified and cross-registered for longitudinal recording by visual inspection (Kitamura et al., 2017). The corresponding Ca^2+^ activity was extracted by applying PCA/ICA analysis in Mosaic software (Inscopix). In total, we recorded 179 BLA *Ppp1r1b*^+^ cells from 5 Ppp1r1b-Cre mice and 169 BLA *Rspo2*^+^ cells from 4 Rspo2-Cre mice.

#### Data analysis

Ca^2+^ events were detected with a threshold 15% ΔF/F. We calculated the average time of Ca^2+^ events, i.e., the sum of all Ca^2+^ events duration/recording time. To identify cells with increased or decreased activity in response to footshocks during CFC on Day 1, we compared the mean ΔF/F of 3 min after delivery of footshocks with that of 3 min habituation stage on Day1. To identify cells with changed neuronal activities during fear retrieval, we compared the mean ΔF/F of the fear retrieval stage (0-3 min) on Day 2 with that of the 3 min habituate period before footshocks on Day 1. To identify cells with increased or decreased activity during fear extinction training, we compared the mean ΔF/F of extinction training with that of fear retrieval stage (0-3 min) (*12*). To identify cells with increased or decreased activity during extinction retrieval, we compared the mean ΔF/F of 5 min extinction retrieval on Day 3 with that of the 3 min fear retrieval stage on Day 2. For each comparison, after shuffling all individual Ca^2+^ transients 5000 times between the two recording sessions to be compared, a distribution of ΔF/F difference was generated. If the actual difference between mean ΔF/F of two recording sessions was higher than 97.5% of the distribution, then the cell was considered as significantly activated (Up cells). If the actual difference between mean ΔF/F of two recording sessions was lower than 2.5% of the distribution, then the cell was considered as significantly inhibited (Down cells). The others were considered as no change cells. All the data were analyzed using custom code written in MATLAB (Mathworks)

#### Cell counting

Images of smFISH counting were acquired using a Zeiss AxioImager.Z2 under a 10X objective. Consecutive sections with 0.2 mm interval (anterior-posterior distance from Bregma, −0.8 mm to −2.6 mm) were used for quantification. Neurons stained for *Fos* (red) and against gene marker (green), *Rspo2* or *Ppp1r1b*, were counted using image analysis software (HALO, Indica Labs), which was compatible with RNAscope Fluorescent Multiplex Kit.

Images of immunohistochemistry counting were acquired using confocal microscopy (Zeiss LSM700) under a 10X objective. Maximum intensity projections were generated using ZEN Blue software (Zeiss). Neurons stained against GFP (green), c-Fos (red), mCherry (red), PP1R1B (green or red) and DAPI (blue) were automatically using cell counting software (ImageJ) with manual adjustment of detection thresholds.

In Figure 6, For DAPI counts, thresholds were set to only include larger nuclei and to exclude non-neuronal small nuclei as previously described. The region of interest (ROI) was defined manually based on anatomical landmarks. All counting was performed blind as to the group and condition that the specimen belonged to.

#### Optogenetic activation/inhibition

To apply optogenetic activation/inhibition during fear extinction in Figure 3, mice were put back to conditioned behavior chamber to receive 45 min fear extinction training. After the first 5 min, the mice received 8 cycles of 3 min optogenetic activation (450-470 nm, 8-12 mW, 20 Hz) or 3 min optical inhibition (520-550 nm, 8-12 mW, constant) with 2 min interval. 450 nm and 520 nm lasers were generated with LED drivers (Doric).

In Figure. 1J-IM, AAV9-DIO-ArchT-EYFP virus was injected into pBLA of Ppp1r1b-Cre mice or Ppp1r1b-Cre-negative mice as control. During fear extinction retrieval on Day 3, the mice received 3 cycles of 1 min optical inhibition (520-550 nm, 8-12 mW, constant) with 1 min interval.

#### Labeling and optogenetic stimulation of fear extinction engram (Related to Figure 4 and 5)

To label fear extinction engram neurons in Figure 4, we injected AAV_9_-c-Fos-tTA together with AAV_9_-TRE-ChR2-EYFP or AAV_9_-TRE-EYFP into the pBLA of C57BL/6 mice, and subjected them to a series of engram labeling and behavioral steps (Figure 4D). This genetic manipulation permits expression of ChR2-EYFP or EYFP in the cells that were activated during fear extinction memory retrieval through the transcriptional promoter of c-Fos gene when doxycycline (Dox) was removed from the diet. Mice were kept on Dox diet after surgery. In Ext-ChR2 group and Ext-EYFP group, mice were switched to Dox-OFF diet for 24 hrs following fear extinction training on Day 2. 60 min after extinction retrieval test on Day 3, mice returned to Dox-ON diet and stayed at home cage on Day 4. In HC-ChR2 group, mice stayed on Dox diet throughout the first round of contextual fear extinction protocol and were switched to Dox-OFF diet for 24 hrs and stayed at home cage on Day 4. On Day 5, mice in all three groups received one immediate footshock (0.75 mA) in the original conditioned context for fear reinstatement. On Day 6, mice received opto-extinction protocol, in which 8 cycles of 3 min-long blue lasers (470 nm, 20 Hz, 8-12 mW) were delivered after the first 5 min. On Day 7, mice were put back to the behavior context for 5 min retrieval test.

In Figure 5, fear extinction engram neurons were labeled in the same fashion as Figure 4, except that the fear extinction engram neurons expressed ArchT-mCherry or mCherry. On Day 4, mice were put back to the conditioned context for 5 min opto-extinction retrieval test, in which mice received three green laser stimulation (520-550 nm, 8-12 mW, constant) at 0-1 min, 2-3 min and 4-5 min time points. For Fos quantification experiment in Figure 5E-5G, mice were sacrificed and the brain were harvested and subjected to smFISH procedure.

#### Labeling and reactivation of reward responsive neurons (Related to Figure 6B, 7B-7G)

To label reward responsive neurons in Figure 6B, One-week post-surgery, mice were deprived of water for 24 hrs while on Dox diet. Water-deprived mice were given regular chaw food without Dox (Dox-OFF diet) for 24 hrs, during which water was given as a reward. 60 min after drinking water, subjects returned to Dox-ON diet to close the labeling window.

After being back to Dox-ON diet, mice were divided into four different groups. Mice in the W/W (Water/Water) group remained on water deprivation for subsequent three days. On Day 5, W/W group mice received *ad libitum* water as a reward 90 min before perfusion. Mice in the W/F (Water/Food) group were deprived of food for subsequent three days and received *ad libitum* food as a reward 90 min before perfusion. During food deprivation, mice received water that contained 200 ul/L Doxycycline to remain on Dox. Mice in the W/Ext (Water/Extinction) group went through three-day contextual fear extinction protocol and were perfused 90 min after extinction retrieval on Day 5. Mice in the W/NonExt group (Water/NonExtinction) group went through the same procedure as the W/Ext group except that they stayed in home cage (HC) instead of extinction training on Day 4. After perfusion, brains of these four groups of mice were sliced and stained for c-Fos staining followed by quantification.

In Figure 7F, water reward responsive neurons were labeled with EYFP or ChR2-EYFP depending on the virus injection combination. Mice went through three-day contextual fear extinction protocol. For EYFP-ON group and ChR2-ON group, mice received opto-extinction protocol, in which 8 cycles of 3 min-long blue lasers (470 nm, 20 Hz, 8-12 mW) were delivered after the first 5 min of extinction training. For ChR2-OFF group, mice didn’t receive any optogenetic stimulation during extinction training.

#### Optogenetic self-stimulation test (Related to Figure 7B and 7C)

The mice were water restricted overnight before the experiment. The optogenetic self-stimulation test was conducted over 60 min session on the following day and no prior training was required. The water-restricted mice were placed in an operant conditioning chamber (Med Associates) equipped with two nose ports (*16, 18*). At the beginning of each trial, each nose port contained 0.05 ml sugar water to initiate the nose poke. No additional water reward was given throughout the trial. One of the two nose ports was randomly assigned as active port, which could deliver 5 s duration of optical stimulation (8-12 mW, 20 Hz, 473 nm) upon nose poke, while the other one did not (inactive). The optical laser was generated through a 470 nm LED light source (XLED1, Lumen Dynamics). The number of nose pokes of active port and inactive port were recorded with Med-PC IV software during 60 min test session. If the difference between active pokes and inactive pokes, (active-inactive), was larger than 20 pokes, it was counted as self-stimulation behavior.

#### Optogenetic place preference (Related to Figures 7B, 7D and 7E)

Mice were place in a custom-made behavioral arena (L45 ×W15.5 × H30 cm, white plexiglass), where each side of the box contained distinct wall cues. Each trial was 20 min and one side was randomly assigned as the stimulation side for each trial. Mice were place in the non-stimulated side at the onset of the experiment. Each time the mouse crossed to the stimulation side of the chamber, mice received continuous 20 Hz laser stimulation (8-12 mW, 473 nm) until the mouse crossed back into the non-stimulation side. Behavioral data were recorded and analyzed with EthoVision XT video tracking software (Noldus Information Technologies).

#### Systemic injection of Kainic acid (Related to Figure S4A and S4B)

To induce seizure, mice were injected intraperitoneally with 15 mg kg^−1^ kainic acid (KA, Tocris Cat. No 0222) after being on Dox off diet for 24 hrs. 3 hrs after KA injection, mice were returned to Dox diet and perfused next day for immunohistochemistry staining.

#### Statistical Analysis

GraphPad Prism (version 7.0c for Mac OS X, GraphPad Software, La Jolla California USA) was used for statistical analysis. All data are presented as mean ± SEM. *n* indicates number of animals or number of cells. Comparisons between two-group data were analyzed by two-tailed unpaired t-test or two-tailed paired t-test. Multiple group comparisons were assessed using a One-way ANOVA or Two-way RM (Repeated Measures) ANOVA, followed by the Tukey’s multiple comparison test when significant main effects or interactions were detected. The null hypothesis was rejected at the *P* < 0.05 level. Data met assumptions of statistical tests. Sample sizes were chosen on the basis of previous studies (Kim et al., 2016; Kim et al., 2017; Liu et al., 2012; Nishi et al., 1997).

**Figure S1.**
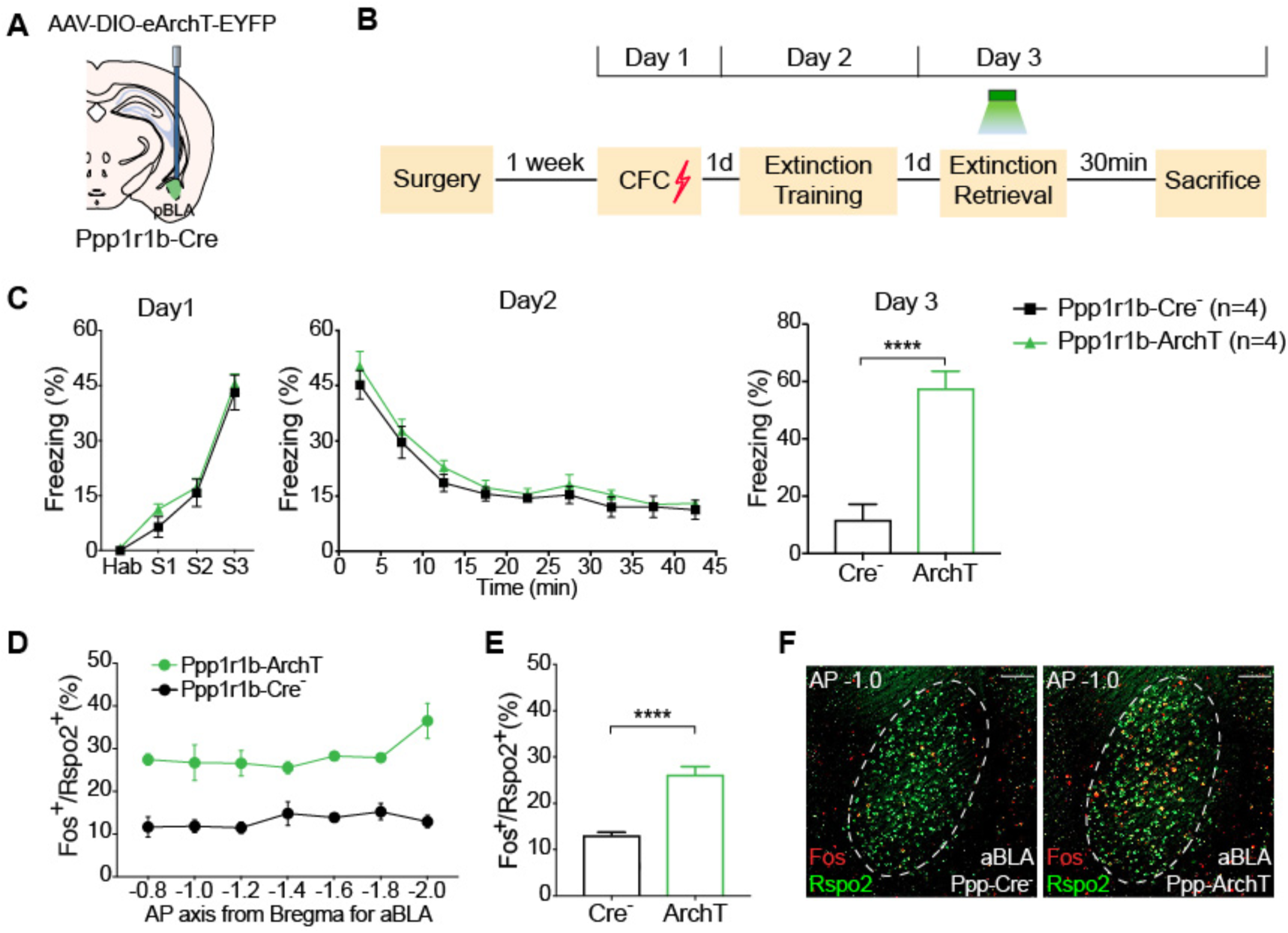
Inhibition of BLA *Ppp1r1b*^+^ neurons during extinction retrieval leads to increased *Rspo2*^+^ neuronal activity (Related to Figure 1) (**A**) Schematic diagram of bilateral injection of AAV_9_-DIO-eArchT-EYFP and optical fiber implant in pBLA of Ppp1r1b-Cre mice or Ppp1r1b-Cre^−^ control mice. (**B**) Experimental protocol. BLA *Ppp1r1b*^+^ neurons were inhibited during extinction retrieval on Day 3 and mice were sacrificed 30 min after extinction retrieval test for smFISH staining. (**C**) Inhibition of *Ppp1r1b*^+^ neurons led to increased freezing level during extinction retrieval on Day 3. Ppp1r1b-Cre- and Ppp1r1b-ArchT mice exhibited similar freezing levels during CFC on Day 1 and extinction training on Day 2. Unpaired t-test. Ppp-Cre^−^ *n* = 4; Ppp-ArchT *n* = 4. (**D**) When BLA *Ppp1r1b*^+^ neurons were inhibited during extinction retrieval on Day 3, percentage of *Fos*^+^ neurons within BLA *Rspo2*^+^ neurons across A/P axis, −0.8 mm to −2.0 mm. (**E**) Average of the percentage of *Fos*^+^/*Rspo2*^+^ shown in (D). Unpaired t-test. (**F**) Double smFISH of Fos (red) and Rspo2 (green) in aBLA in Ppp-Cre^−^ mice (left) and Ppp-ArchT mice (right). \**P* < 0.05, \*\**P* < 0.01, \*\*\**P* < 0.001, \*\*\*\**P* < 0.0001. Data are presented as mean ± SEM. Scale bars: 200 µm (M)

**Figure S2.**
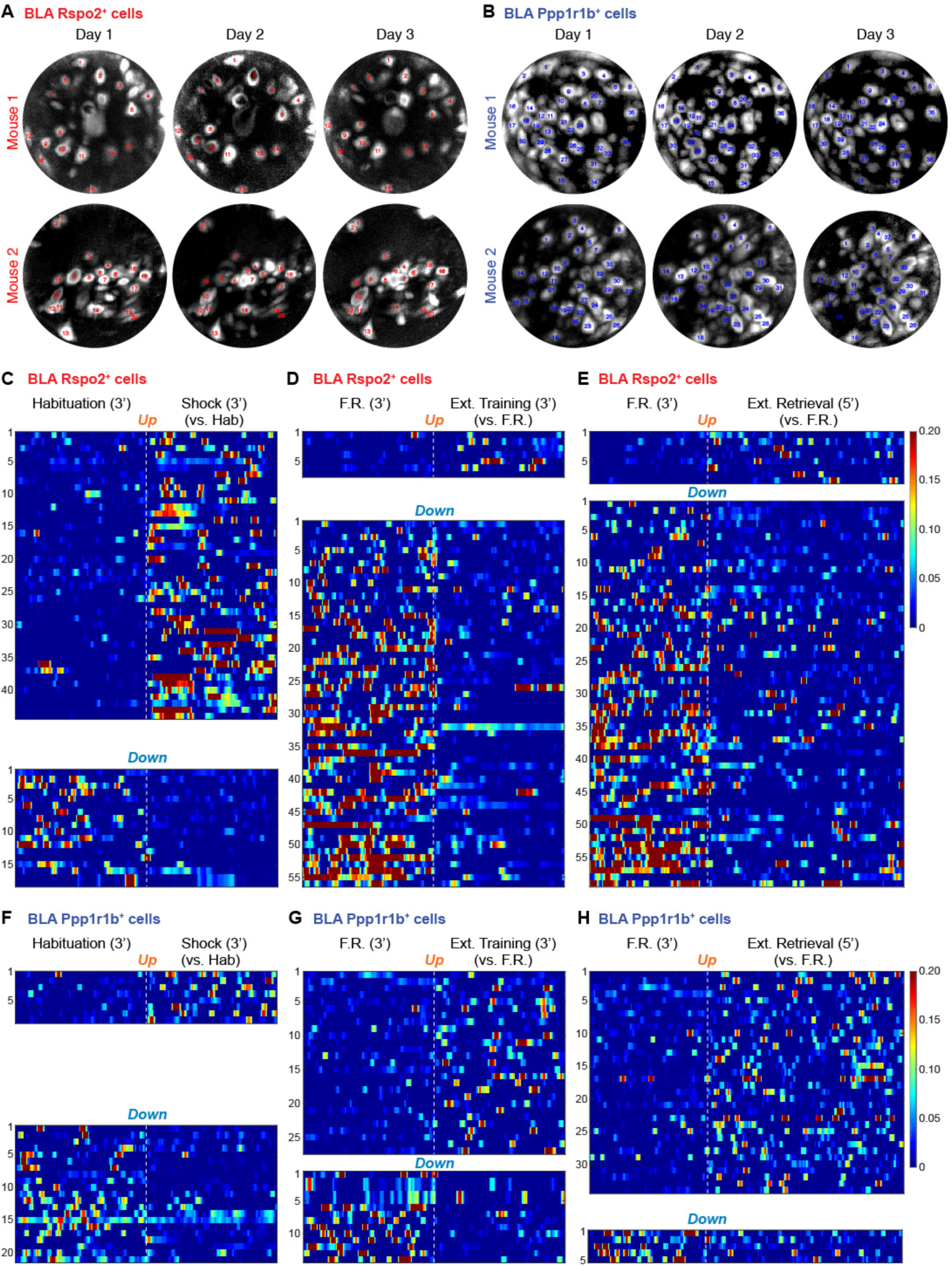
Longitudinal recording of BLA *Rspo2*^+^ and *Ppp1r1b*^+^ cells (Related to Figure 2) (**A**) Stacked filed of view (FOV) images of BLA *Rspo2*^+^ neurons acquired on Day 1 CFC, Day 2 Extinction training and Day 3 Extinction retrieval. Same cell numbers within each mouse are tracked across three days. (**B**) Stacked FOV images of BLA *Ppp1r1b*^+^ neurons acquired on Day 1 CFC, Day 2-Extinction training and Day 3-Extinction retrieval. Same cell numbers within each mouse are tracked across three days. (**C**) Heat maps of Ca^2+^ responses of BLA *Rspo2*^+^ neurons that were activated (top-Up, *n* = 44) and inhibited (bottom-Down, *n* = 18) by shocks during CFC on Day 1. (**D**) Heat maps of Ca^2+^ responses of BLA *Rspo2*^+^ neurons that were activated (top-Up, *n* = 7) and inhibited (bottom-Down, *n* = 56) during extinction training. Extinction training shows the recording of 35-38 min timestamp on Day 2. (**E**) Heat map of Ca^2+^ responses of BLA *Rspo2*^+^ neurons that were activated (top-Up, *n* = 8) and inhibited (bottom-Down, *n* = 59) during extinction retrieval. (**F**) Heat map of Ca^2+^ responses of BLA *Ppp1r1b*^+^ neurons that were activated (top-Up, *n* = 8) and inhibited (bottom-Down, *n* = 21) after shocks during CFC on Day 1. (**G**) Heat map of Ca^2+^ responses of BLA *Ppp1r1b*^+^ neurons that were activated (top-Up, *n* = 28) or inhibited (bottom-Down, *n* = 14) during extinction training. Extinction training shows the recording of 35-38 min timestamp on Day 2. (**H**) Heat map of Ca^2+^ responses of BLA *Ppp1r1b*^+^ neurons that were activated (top-Up, *n* = 34) or inhibited (bottom-Down, *n* = 5) during extinction retrieval.

**Figure S3.**
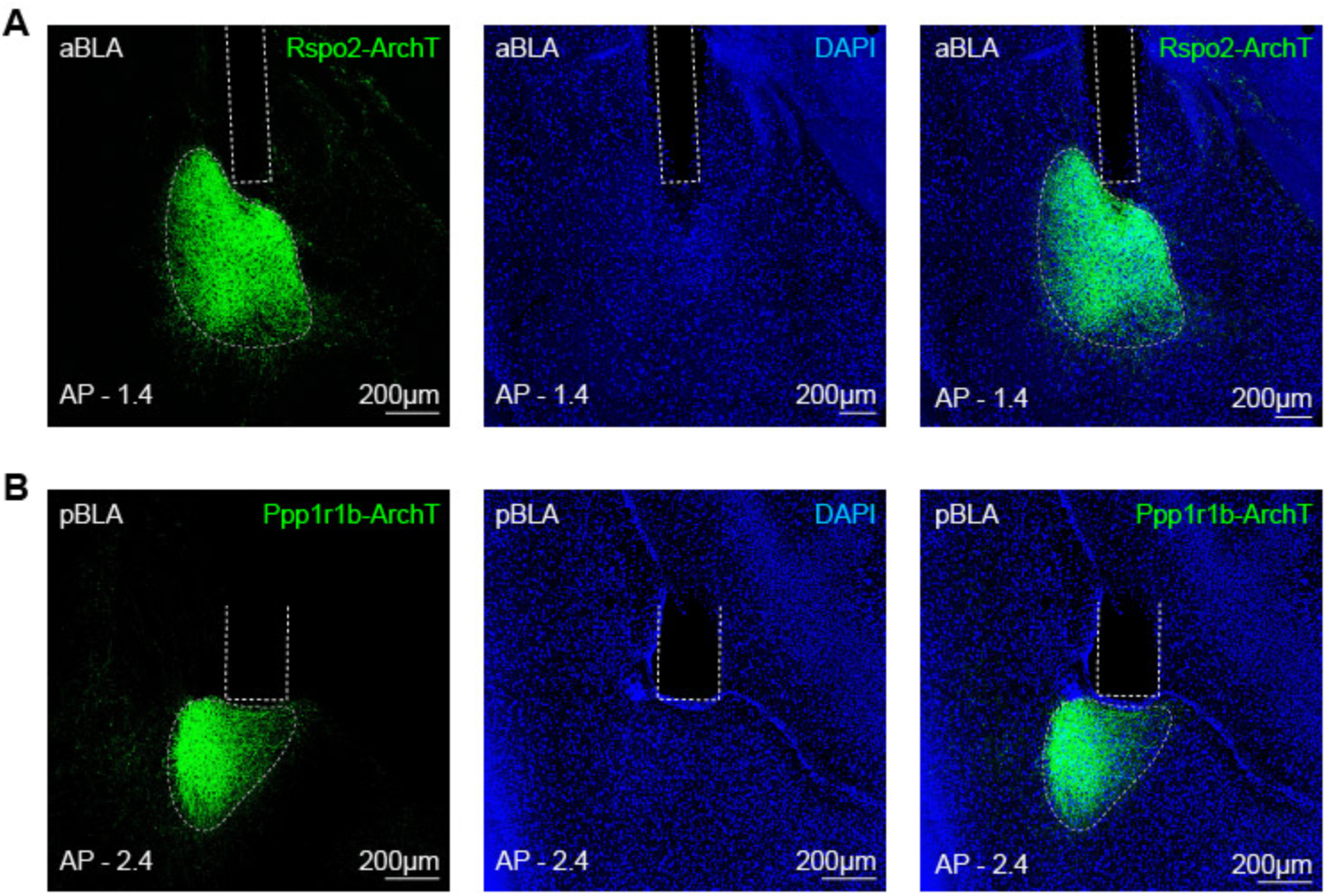
Expression of eArchT-EYFP and fiber placement for targeting BLA *Rspo2*^+^ and *Ppp1r1b*^+^ neurons (Related to Figure 3) (**A**) Representative histology images showing the expression of eArchT-EYFP and optical fiber implant in Rspo2-Cre mice. (**B**) Representative histology images showing the expression of eArchT-EYFP and optical fiber implant in Ppp1r1b-Cre mice.

**Figure S4.**
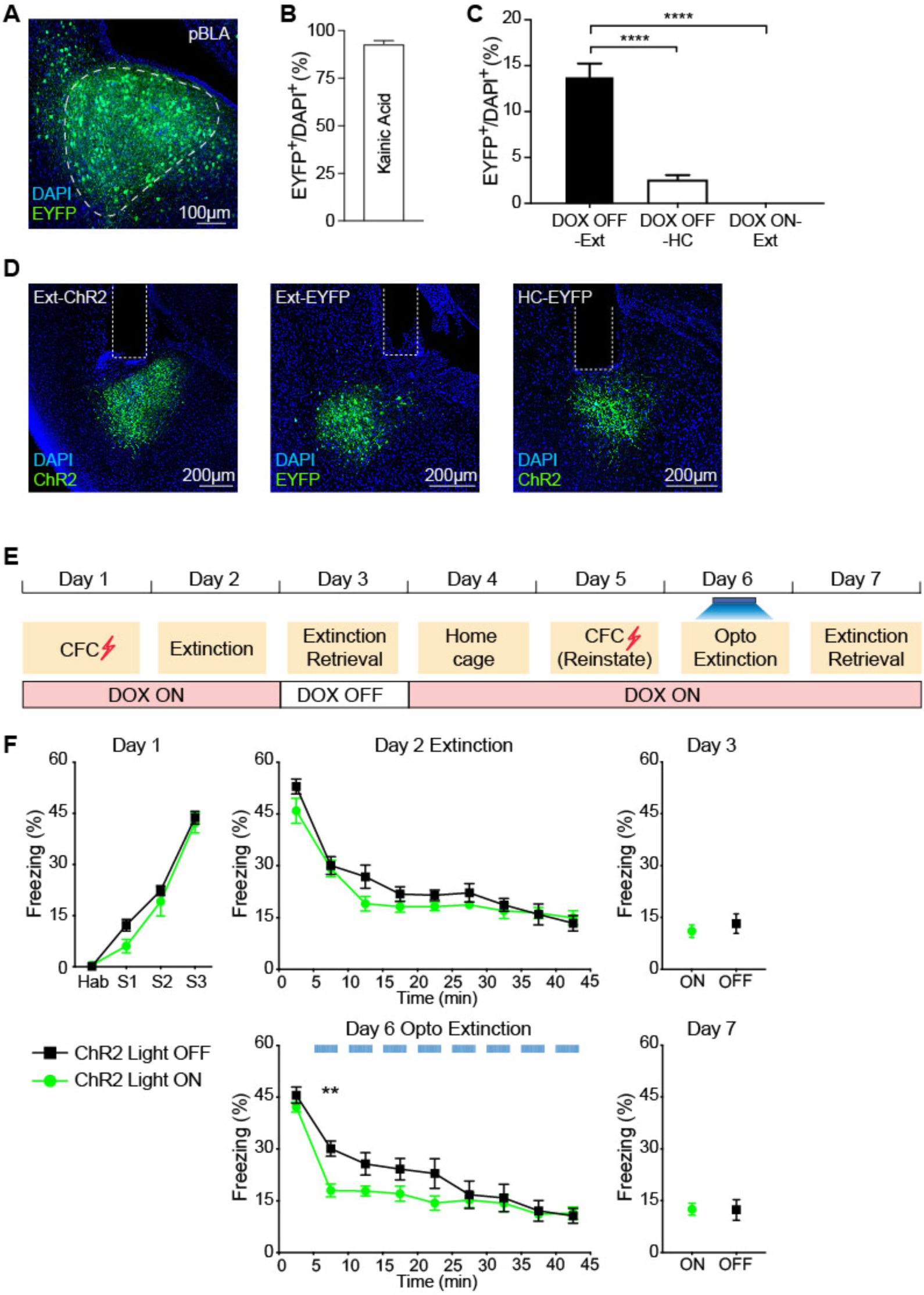
Characterization of fear extinction engram cells in pBLA (Related to Figure 4) (**A**) AAV_9_-c-fos-tTA and AAV_9_-TRE-EYFP virus combination were injected into pBLA of C57BL/6 mice. After 24 hrs Dox-OFF diet, mice were injected with Kainic Acid to induce seizure. Image showing efficient labeling in pBLA. (**B**) EYFP^+^ cell counts that were activated by Kainic Acid-induced seizure from pBLA section (*n* = 6 mice). 93 ± 3% cells were labelled. (**C**) EYFP^+^ cell counts from pBLA sections under three different conditions: Dox OFF during extinction retrieval (Dox OFF-Ext, *n* = 4 mice), Dox OFF at home cage (Dox OFF-HC, *n* = 4 mice) and Dox ON during extinction retrieval (Dox ON-Ext, *n* = 4 mice). One-way ANOVA. (**D**) Representative histology image showing expression of ChR2-EYFP and EYFP and optical fiber implants. (**E**) Behavior paradigm as described in Figure 4D. (**F**) When ChR2-expressing extinction engram neurons were activated, Light ON group exhibited accelerated fear extinction learning on Day 6 compared to light OFF group. ChR2 Light OFF, *n* = 7, ChR2 Light ON, *n* = 12. Two-way RM ANOVA. \**P* < 0.05, \*\**P* < 0.01, \*\*\**P* < 0.001, \*\*\*\**P* < 0.0001. Data are presented as mean ± SEM.

**Figure S5.**
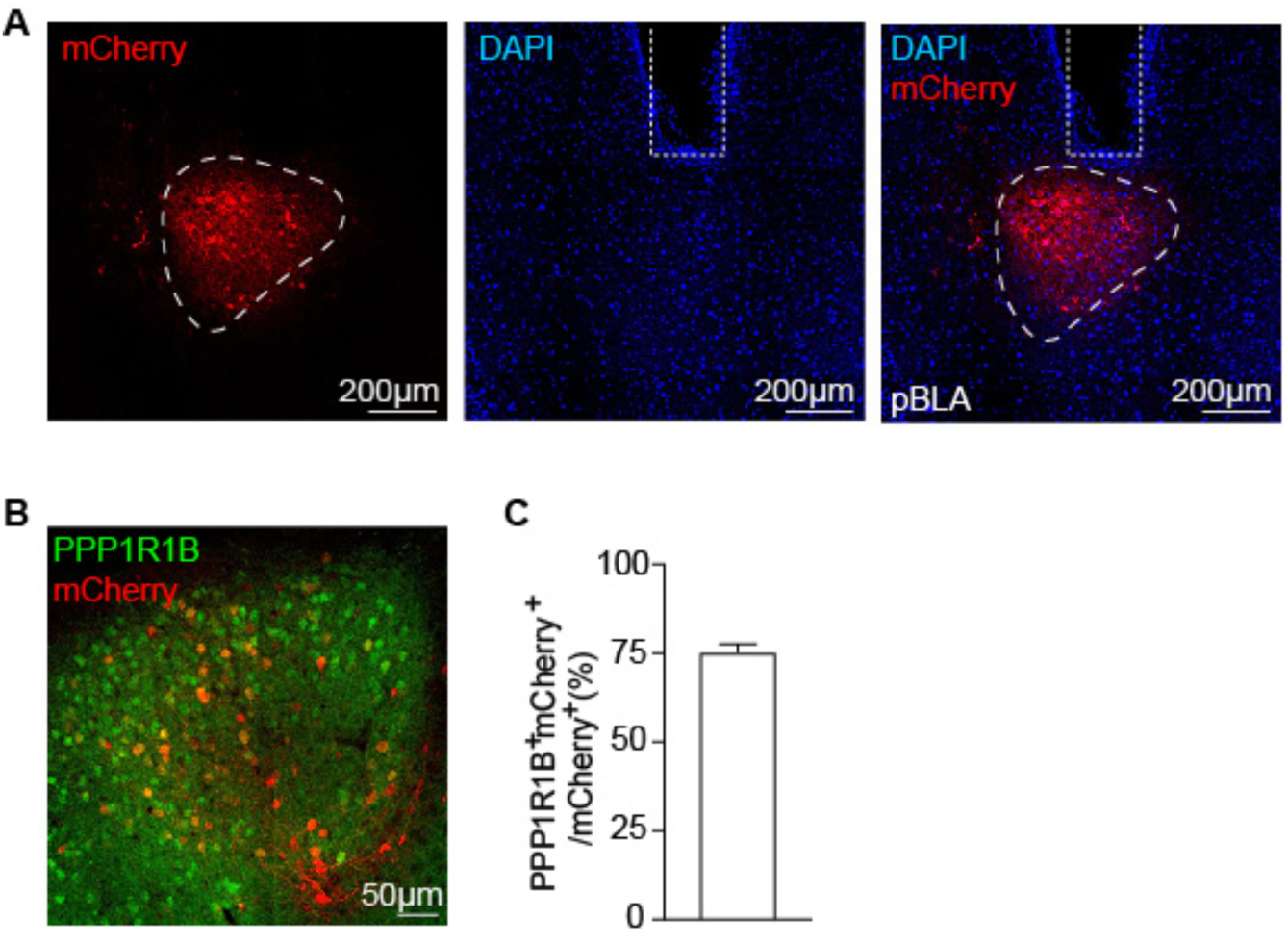
Inhibition of fear extinction engram in pBLA (Related to Figure 5) (**A**) AAV_9_-c-fos-tTA and AAV_9_-TRE-ArchT-mCherry virus combination were injected into pBLA of C57BL/6 mice. Mice underwent fear extinction retrieval during 24 hrs Dox OFF period. Image showing efficient labeling of eArchT-mCherry in pBLA. (**B**) Image showing activity-induced mCherry-expressing neurons in pBLA stained with PPP1R1B antibody (green). (**C**) 75 ± 2% mCherry^+^ cells were labeled as PPP1R1B^+^. Data are presented as mean ± SEM.

**Figure S6.**
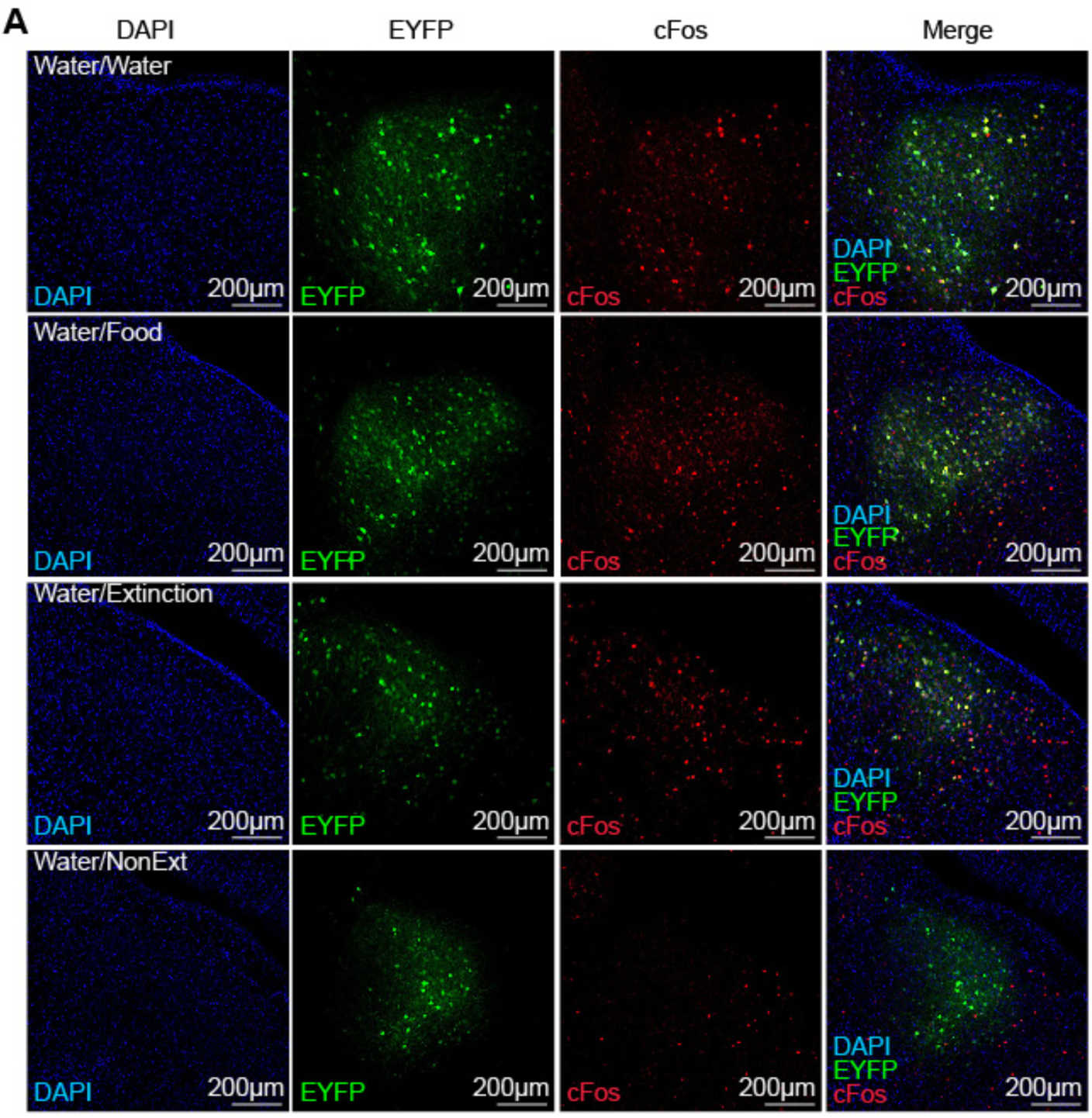
Cellular overlap of water-reward neurons and extinction engram neurons in pBLA (Related to Figure 6) (**A**) pBLA neurons activated by water reward was labeled by EYFP (green) and pBLA cells reactivated by water reward (Water/Water group), food reward (Water/Food group), extinction retrieval (Water/Extinction group) and fear retrieval (Water/NonExtinction group) were stained by c-Fos (red). The merged images with DAPI staining are shown on the right.

**Figure S7.**
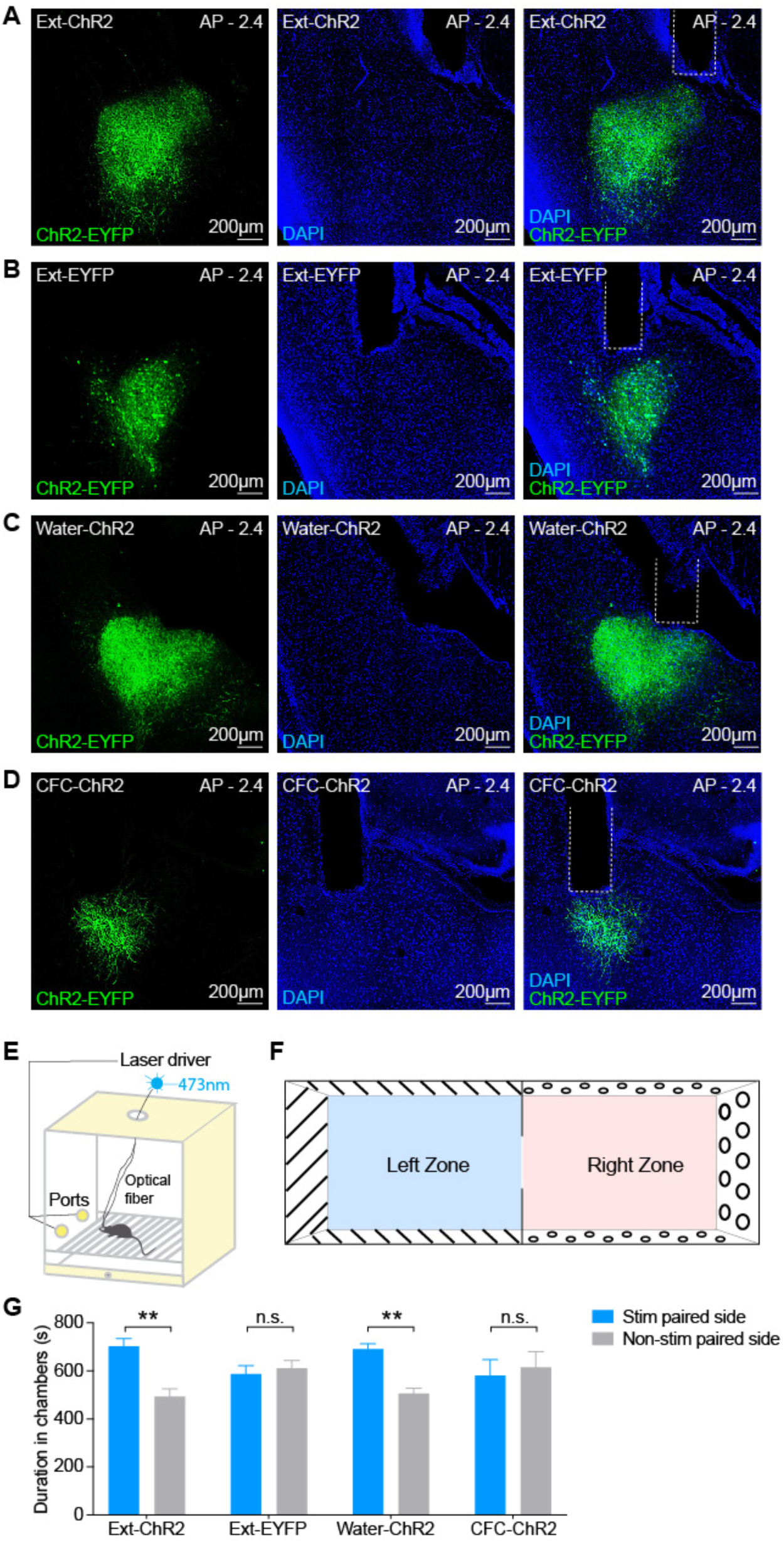
Behavioral overlap of water-reward neurons and extinction engram neurons in pBLA (Related to Figure 7) (**A**) Image showing efficient labeling of ChR2-EYFP and optical fiber implant in pBLA of Ext-ChR2 group in Figure 7B. (**B**) Image showing efficient labeling of EYFP and optical fiber implant in pBLA of Ext-EYFP group in Figure 7B. (**C**) Image showing efficient labeling of ChR2-EYFP and optical fiber implant in pBLA of Water-ChR2 group in Figure 7B. (**D**) Image showing labeling of ChR2-EYFP and optical fiber implant in pBLA of CFC-ChR2 group in Figure 7B. (**E**) Schematic diagram of operant chamber for optogenetic self-stimulation behavior in Figure 7C. (**F**) Schematic diagram of chamber used for optogenetic place preference behavior in Figure 7D. (**G**) Total duration spent in the stimulation paired side chamber and non-stimulation paired side. Ext-ChR2 and Water-ChR2 group spent significantly more time in the stimulation paired side chamber but not Ext-EYFP and CFC-ChR2 group. Paired t-test. \**P* < 0.05, \*\**P* < 0.01, \*\*\**P* < 0.001, \*\*\*\**P* < 0.0001. Data are presented as mean ± SEM.

