## Supplementary material for "Amygdala Reward Neurons Form and Store Fear Extinction Memory"

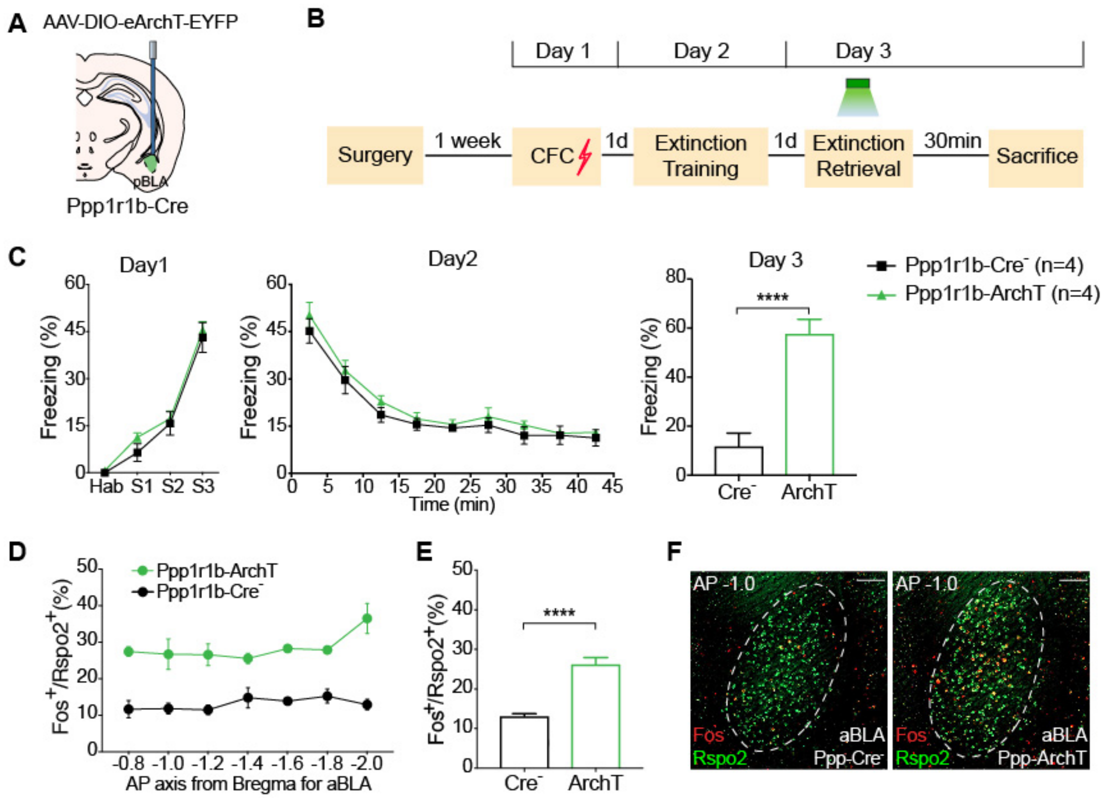
A
AAV-DIO-eArchT-EYFP
pBLA
Ppp1r1b-Cre
B
Day 1
Day 2
Day 3
Surgery
1 week
CFC
1d
Extinction Training
1d
Extinction Retrieval
30min
Sacrifice
C
Day1
Freezing (%)
Hab S1 S2 S3
Day2
Time (min)
Day 3
****
Cre⁻
ArchT
Ppp1r1b-Cre⁻ (n=4)
Ppp1r1b-ArchT (n=4)
D
Ppp1r1b-ArchT
Ppp1r1b-Cre⁻
Fos⁺/Rspo2⁺(%)
AP axis from Bregma for aBLA
E
Fos⁺/Rspo2⁺(%)
****
Cre⁻
ArchT
F
AP -1.0
Fos
Rspo2
aBLA
Ppp-Cre⁻
AP -1.0
Fos
Rspo2
aBLA
Ppp-ArchT

1

**Figure S1. Inhibition of BLA *Ppp1r1b*$^+$ neurons during extinction retrieval leads to increased *Rspo2*$^+$ neuronal activity (Related to Figure 1)**

(**A**) Schematic diagram of bilateral injection of $AAV_9$-DIO-eArchT-EYFP and optical fiber implant in pBLA of Ppp1r1b-Cre mice or Ppp1r1b-Cre$^-$ control mice. (**B**) Experimental protocol. BLA *Ppp1r1b*$^+$ neurons were inhibited during extinction retrieval on Day 3 and mice were sacrificed 30 min after extinction retrieval test for smFISH staining. (**C**) Inhibition of *Ppp1r1b*$^+$ neurons led to increased freezing level during extinction retrieval on Day 3. Ppp1r1b-Cre- and Ppp1r1b-ArchT mice exhibited similar freezing levels during CFC on Day 1 and extinction training on Day 2. Unpaired t-test. Ppp-Cre$^-$ $n$ = 4; Ppp-ArchT $n$ = 4. (**D**) When BLA *Ppp1r1b*$^+$ neurons were inhibited during extinction retrieval on Day 3, percentage of *Fos*$^+$ neurons within BLA *Rspo2*$^+$ neurons across A/P axis, -0.8 mm to -2.0 mm. (**E**) Average of the percentage of *Fos*$^+$/*Rspo2*$^+$ shown in (D). Unpaired t-test. (**F**) Double smFISH of Fos (red) and Rspo2 (green) in aBLA in Ppp-Cre$^-$ mice (left) and Ppp-ArchT mice (right). *$P < 0.05$, **$P < 0.01$, ***$P < 0.001$, ****$P < 0.0001$. Data are presented as mean ± SEM. Scale bars: 200 μm (M)

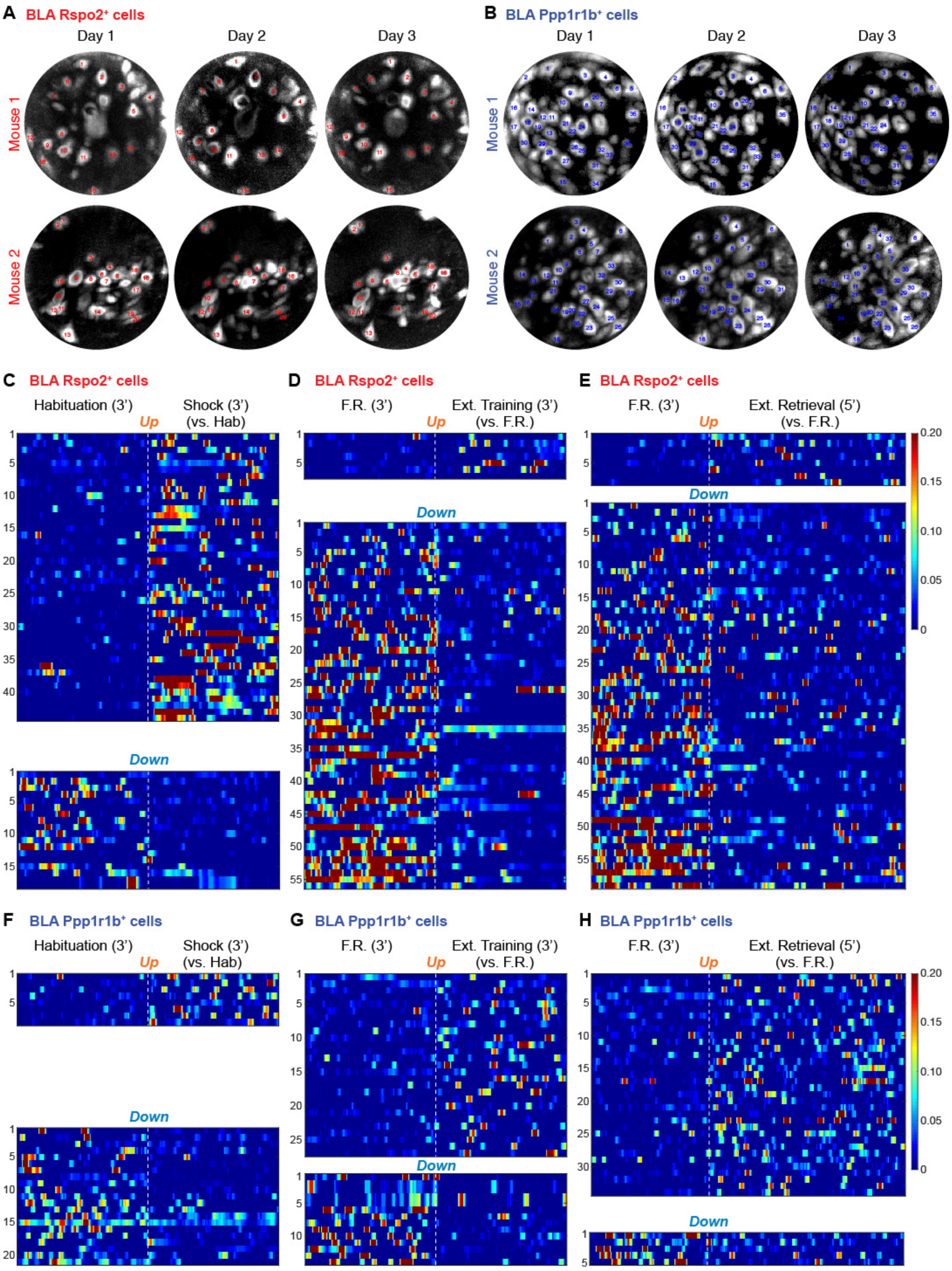
A BLA Rspo2+ cells
Day 1
Day 2
Day 3
Mouse 1
Mouse 2
B BLA Ppp1r1b+ cells
Day 1
Day 2
Day 3
Mouse 1
Mouse 2
C BLA Rspo2+ cells
Habituation (3')
Shock (3') (vs. Hab)
Up
Down
D BLA Rspo2+ cells
F.R. (3')
Ext. Training (3') (vs. F.R.)
Up
Down
E BLA Rspo2+ cells
F.R. (3')
Ext. Retrieval (5') (vs. F.R.)
Up
Down
0.20
0.15
0.10
0.05
0
F BLA Ppp1r1b+ cells
Habituation (3')
Shock (3') (vs. Hab)
Up
Down
G BLA Ppp1r1b+ cells
F.R. (3')
Ext. Training (3') (vs. F.R.)
Up
Down
H BLA Ppp1r1b+ cells
F.R. (3')
Ext. Retrieval (5') (vs. F.R.)
Up
Down
0.20
0.15
0.10
0.05
0

**Figure S2. Longitudinal recording of BLA *Rspo2*$^+$ and *Ppp1r1b*$^+$ cells (Related to Figure 2)**

(**A**) Stacked filed of view (FOV) images of BLA *Rspo2*$^+$ neurons acquired on Day 1 CFC, Day 2 Extinction training and Day 3 Extinction retrieval. Same cell numbers within each mouse are tracked across three days. (**B**) Stacked FOV images of BLA *Ppp1r1b*$^+$ neurons acquired on Day 1 CFC, Day 2-Extinction training and Day 3-Extinction retrieval. Same cell numbers within each mouse are tracked across three days. (**C**) Heat maps of $Ca^{2+}$ responses of BLA *Rspo2*$^+$ neurons that were activated (top-Up, $n$ = 44) and inhibited (bottom-Down, $n$ = 18) by shocks during CFC on Day 1. (**D**) Heat maps of $Ca^{2+}$ responses of BLA *Rspo2*$^+$ neurons that were activated (top-Up, $n$ = 7) and inhibited (bottom-Down, $n$ = 56) during extinction training. Extinction training shows the recording of 35-38 min timestamp on Day 2. (**E**) Heat map of $Ca^{2+}$ responses of BLA *Rspo2*$^+$ neurons that were activated (top-Up, $n$ = 8) and inhibited (bottom-Down, $n$ = 59) during extinction retrieval. (**F**) Heat map of $Ca^{2+}$ responses of BLA *Ppp1r1b*$^+$ neurons that were activated (top-Up, $n$ = 8) and inhibited (bottom-Down, $n$ = 21) after shocks during CFC on Day 1. (**G**) Heat map of $Ca^{2+}$ responses of BLA *Ppp1r1b*$^+$ neurons that were activated (top-Up, $n$ = 28) or inhibited (bottom-Down, $n$ = 14) during extinction training. Extinction training shows the recording of 35-38 min timestamp on Day 2. (**H**) Heat map of $Ca^{2+}$ responses of BLA *Ppp1r1b*$^+$ neurons that were activated (top-Up, $n$ = 34) or inhibited (bottom-Down, $n$ = 5) during extinction retrieval.

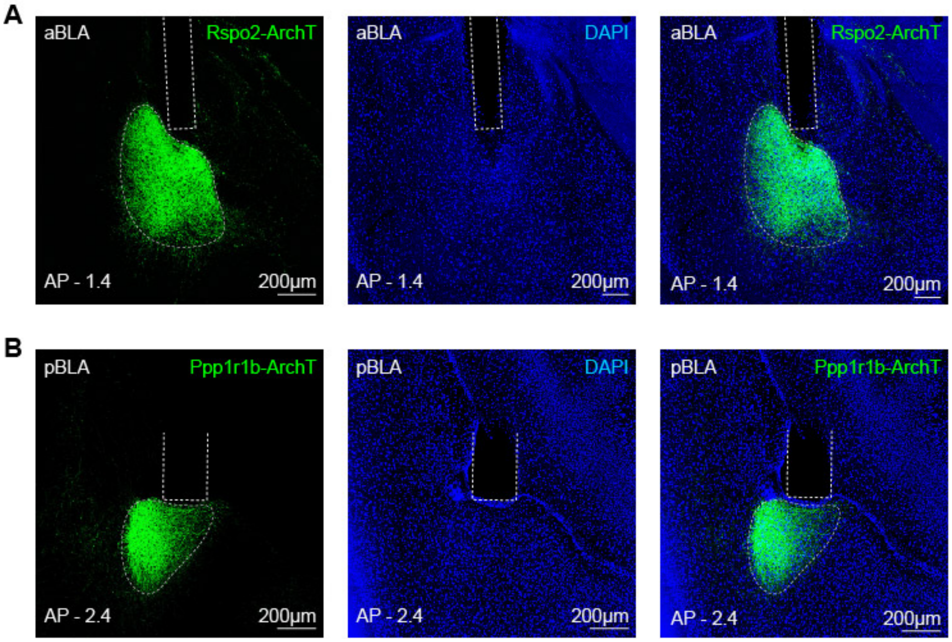


**Figure S3. Expression of eArchT-EYFP and fiber placement for targeting BLA *Rspo2*$^{+}$ and *Ppp1r1b*$^{+}$ neurons (Related to Figure 3)**

(**A**) Representative histology images showing the expression of eArchT-EYFP and optical fiber implant in Rspo2-Cre mice. (**B**) Representative histology images showing the expression of eArchT-EYFP and optical fiber implant in Ppp1r1b-Cre mice.

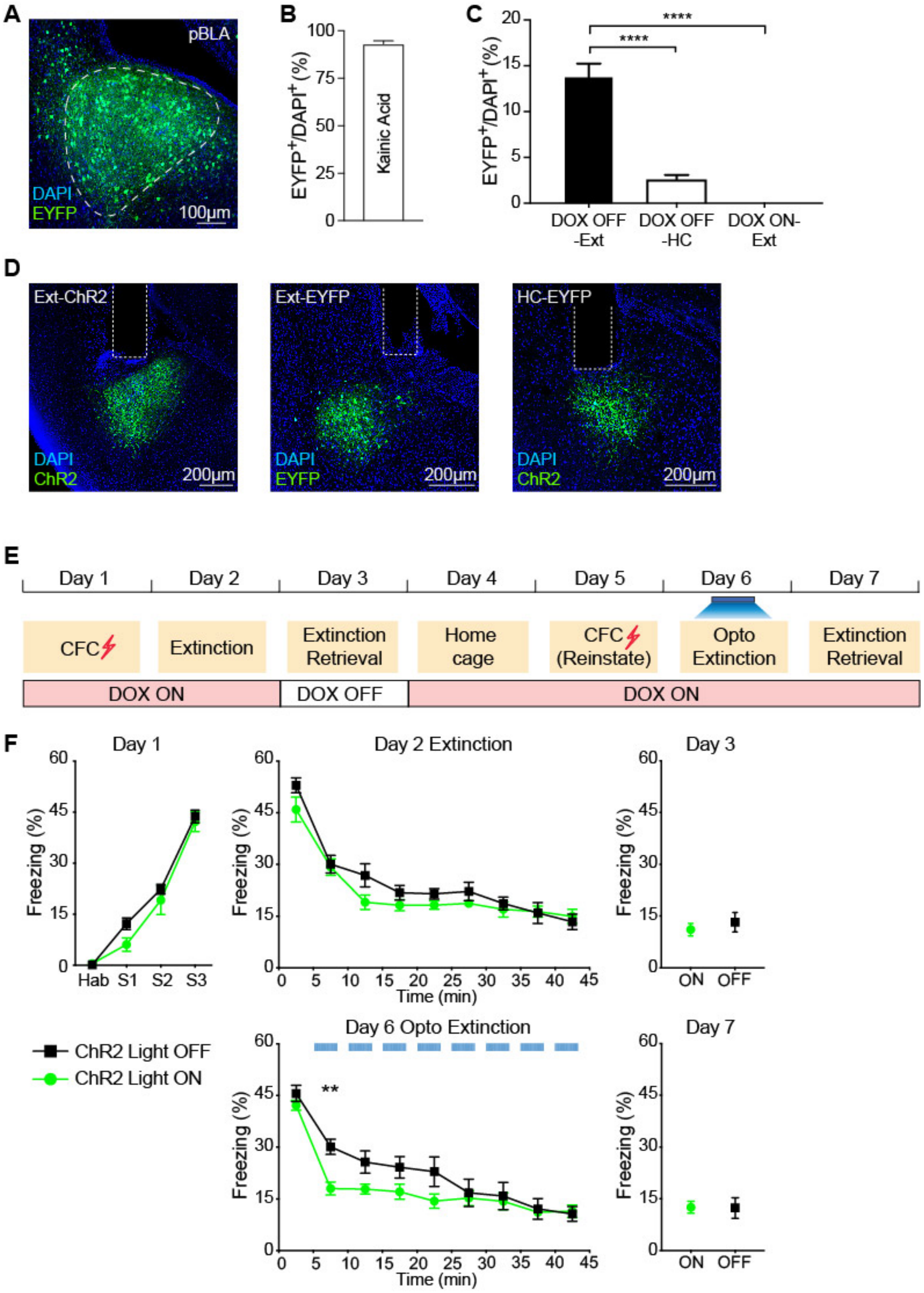
A
pBLA
DAPI
EYFP
100μm
B
EYFP+/DAPI+ (%)
Kainic Acid
C
EYFP+/DAPI+ (%)
****
****
DOX OFF -Ext
DOX OFF -HC
DOX ON- Ext
D
Ext-ChR2
Ext-EYFP
HC-EYFP
DAPI
ChR2
EYFP
200μm
E
Day 1
Day 2
Day 3
Day 4
Day 5
Day 6
Day 7
CFC
Extinction
Extinction Retrieval
Home cage
CFC (Reinstate)
Opto Extinction
Extinction Retrieval
DOX ON
DOX OFF
DOX ON
F
Day 1
Day 2 Extinction
Day 3
Freezing (%)
Hab
S1
S2
S3
Time (min)
ON
OFF
ChR2 Light OFF
ChR2 Light ON
Day 6 Opto Extinction
**
Day 7

**Figure S4. Characterization of fear extinction engram cells in pBLA (Related to Figure 4)**

(**A**) $AAV_9$-c-fos-tTA and $AAV_9$-TRE-EYFP virus combination were injected into pBLA of C57BL/6 mice. After 24 hrs Dox-OFF diet, mice were injected with Kainic Acid to induce seizure. Image showing efficient labeling in pBLA. (**B**) $EYFP^+$ cell counts that were activated by Kainic Acid-induced seizure from pBLA section ($n$ = 6 mice). 93 ± 3% cells were labelled. (**C**) $EYFP^+$ cell counts from pBLA sections under three different conditions: Dox OFF during extinction retrieval (Dox OFF-Ext, $n$ = 4 mice), Dox OFF at home cage (Dox OFF-HC, $n$ = 4 mice) and Dox ON during extinction retrieval (Dox ON-Ext, $n$ = 4 mice). One-way ANOVA. (**D**) Representative histology image showing expression of ChR2-EYFP and EYFP and optical fiber implants. (**E**) Behavior paradigm as described in Figure 4D. (**F**) When ChR2-expressing extinction engram neurons were activated, Light ON group exhibited accelerated fear extinction learning on Day 6 compared to light OFF group. ChR2 Light OFF, $n$ = 7, ChR2 Light ON, $n$ = 12. Two-way RM ANOVA. *$P < 0.05$, **$P < 0.01$, ***$P < 0.001$, ****$P < 0.0001$. Data are presented as mean ± SEM.

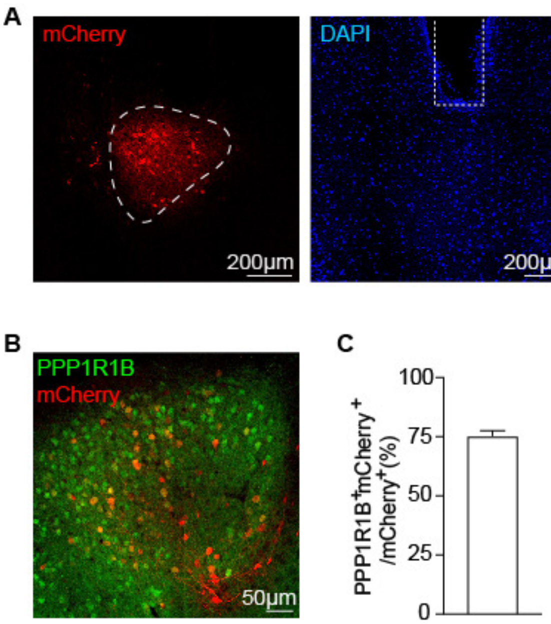


**Figure S5. Inhibition of fear extinction engram in pBLA (Related to Figure 5)**

(**A**) $AAV_9$-c-fos-tTA and $AAV_9$-TRE-ArchT-mCherry virus combination were injected into pBLA of C57BL/6 mice. Mice underwent fear extinction retrieval during 24 hrs Dox OFF period. Image showing efficient labeling of eArchT-mCherry in pBLA. (**B**) Image showing activity-induced mCherry-expressing neurons in pBLA stained with PPP1R1B antibody (green). (**C**) 75 ± 2% mCherry$^+$ cells were labeled as PPP1R1B$^+$. Data are presented as mean ± SEM.

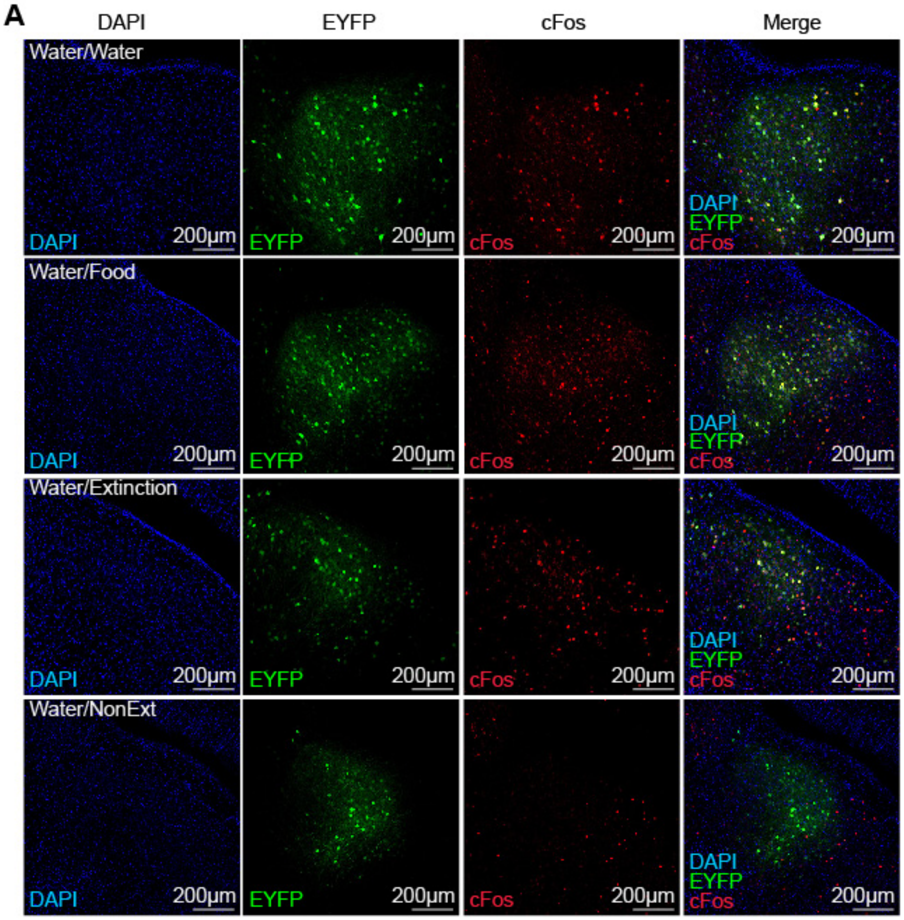
A
DAPI
EYFP
cFos
Merge
Water/Water
Water/Food
Water/Extinction
Water/NonExt
DAPI
EYFP
cFos
200µm

**Figure S6. Cellular overlap of water-reward neurons and extinction engram neurons in pBLA (Related to Figure 6)**

(**A**) pBLA neurons activated by water reward was labeled by EYFP (green) and pBLA cells reactivated by water reward (Water/Water group), food reward (Water/Food group), extinction retrieval (Water/Extinction group) and fear retrieval (Water/NonExtinction group) were stained by c-Fos (red). The merged images with DAPI staining are shown on the right.

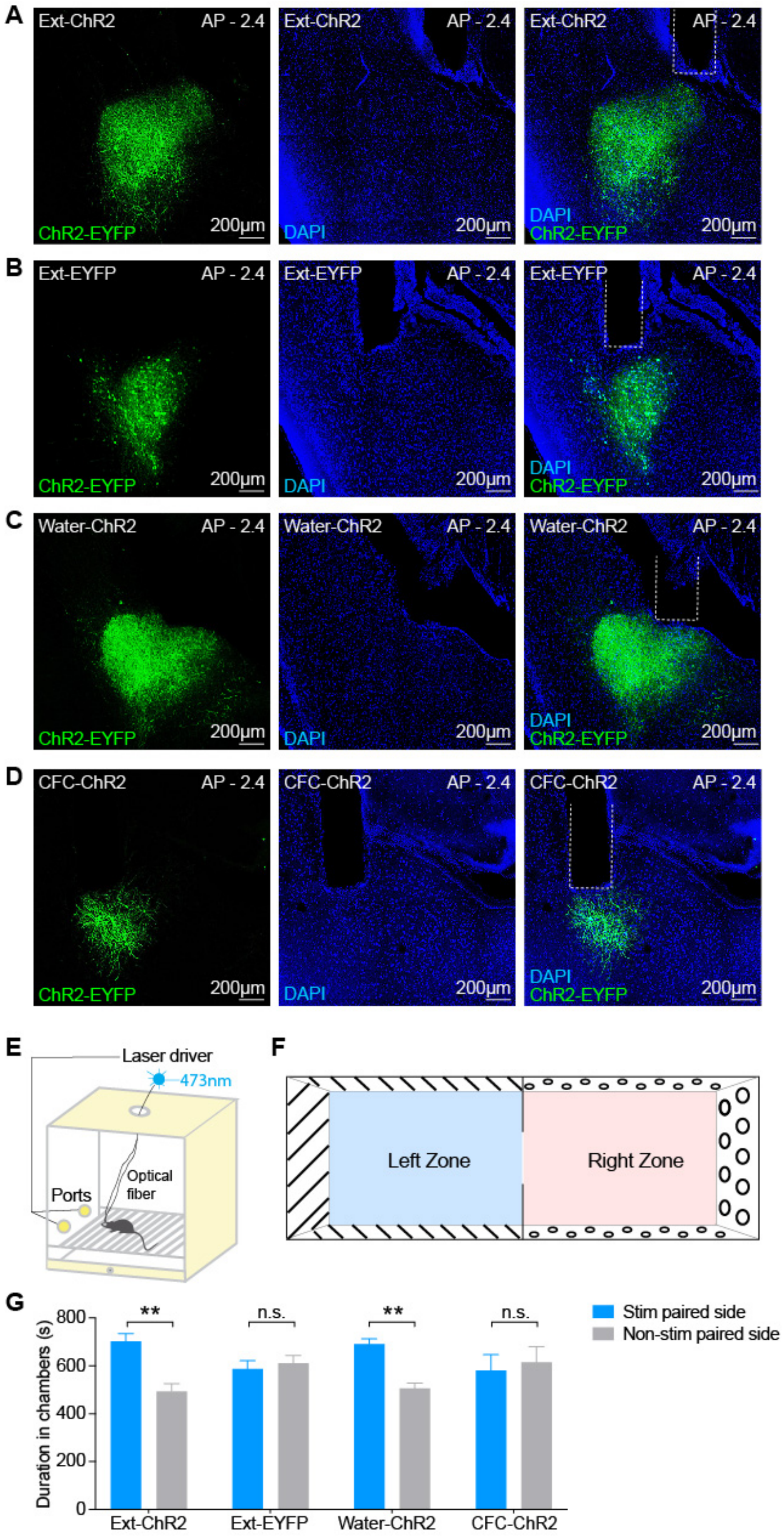
A
Ext-ChR2
AP - 2.4
ChR2-EYFP
200μm
DAPI
B
Ext-EYFP
C
Water-ChR2
D
CFC-ChR2
E
Laser driver
473nm
Ports
Optical fiber
F
Left Zone
Right Zone
G
Duration in chambers (s)
800
600
400
200
0
**
n.s.
Stim paired side
Non-stim paired side
Ext-ChR2
Ext-EYFP
Water-ChR2
CFC-ChR2

1

**Figure S7. Behavioral overlap of water-reward neurons and extinction engram neurons in pBLA (Related to Figure 7)**

(A) Image showing efficient labeling of ChR2-EYFP and optical fiber implant in pBLA of Ext-ChR2 group in Figure 7B. (**B**) Image showing efficient labeling of EYFP and optical fiber implant in pBLA of Ext-EYFP group in Figure 7B. (**C**) Image showing efficient labeling of ChR2-EYFP and optical fiber implant in pBLA of Water-ChR2 group in Figure 7B. (**D**) Image showing labeling of ChR2-EYFP and optical fiber implant in pBLA of CFC-ChR2 group in Figure 7B. (**E**) Schematic diagram of operant chamber for optogenetic self-stimulation behavior in Figure 7C. (**F**) Schematic diagram of chamber used for optogenetic place preference behavior in Figure 7D. (**G**) Total duration spent in the stimulation paired side chamber and non-stimulation paired side. Ext-ChR2 and Water-ChR2 group spent significantly more time in the stimulation paired side chamber but not Ext-EYFP and CFC-ChR2 group. Paired t-test. *$P < 0.05$, **$P < 0.01$, ***$P < 0.001$, ****$P < 0.0001$. Data are presented as mean ± SEM.
